# Contact with the CsrA core is required for allosteric inhibition by FliW in *Bacillus subtilis*

**DOI:** 10.1101/2020.07.02.184341

**Authors:** Reid T. Oshiro, Daniel B. Kearns

## Abstract

The RNA-binding protein CsrA is a post-transcriptional regulator that is encoded in genomes throughout the bacterial phylogeny. In the gamma-proteobacteria, the activity of CsrA is inhibited by small RNAs that competitively sequester CsrA binding. In contrast, the firmicute *Bacillus subtilis* encodes a protein inhibitor of CsrA called FliW, that non-competitively inhibits CsrA activity but the precise mechanism of antagonism is unclear. Here we take an unbiased genetic approach to identify residues of FliW important for CsrA inhibition that fall into two distinct spatial and functional classes. Most loss-of-function alleles mutated FliW residues that surround the critical regulatory CsrA residue N55 and abolished CsrA interaction. Two loss-of-function alleles however mutated FliW residues near the CsrA core dimerization domain and maintained interaction with CsrA. One of these two alleles reversed charge at what appeared to be a salt bridge with the CsrA core region, charge reversal of the CsrA partner residue phenocopied the FliW allele, and charge reversal of both residues simultaneously restored antagonism. We propose a model in which initial interaction between FliW and CsrA is necessary but not sufficient for antagonism which also requires salt bridge formation with, and deformation of, the CsrA core domain to allosterically abolish RNA binding activity.

**Summary:** CsrA is a small dimeric protein that binds RNA and is one of the few known examples of transcript-specific translational regulators in bacteria. A protein called FliW binds to and antagonizes CsrA; despite having a high-resolution three-dimensional structure of the FliW-CsrA complex, the mechanism of non-competitive inhibition remains unresolved. Here we identify FliW residues required for antagonism and we find that the residues make a linear connection in the complex from initial binding interaction with CsrA to a critical salt bridge near the core of the CsrA dimer. We propose that the salt bridge represents an allosteric contact that distorts the CsrA core to prevent RNA binding.

## Introduction

CsrA is a bacterial RNA-binding protein and post-transcriptional regulator (Liu and Romeo, 1997). Primarily an inhibitor of translation, CsrA dimers bind to tandem hairpin structures in a transcript and often occludes the Shine-Dalgarno ribosome binding site (Dubey et al., 2005; Mercante et al., 2006; Schubert et al., 2007; Mercante et al., 2009). In enteric gamma-proteobacteria, CsrA pleiotropically controls translation of a wide range of genes involved in physiological transition during pathogenesis (Vakulskas et al., 2015; Esquerré et al., 2016; Holmqvist 2016; Potts et al., 2017; Potts et al., 2018). Repression is relieved by the expression of small RNAs (sRNA) with multiple CsrA binding sites that act as competitive inhibitors and sequester CsrA (Liu et al., 1997; Suzuki et al., 2002; Weilbacher et al., 2003; Kulkarni et al., 2006; Zere et al., 2015). In the Gram-positive bacterium *Bacillus subtilis* however, CsrA appears dedicated to flagellar regulation and inhibits translation of perhaps only a single gene, flagellin (Yakhnin et al., 2007; Mukherjee et al., 2011; Oshiro et al., 2019). The flagellin transcript is the most abundant transcript in the cell when expressed, and thus would be difficult to outcompete if CsrA was regulated by a competitive mechanism (Mondal et al., 2016). Instead, CsrA is antagonized non-competitively by a stoichiometric ratio of the protein FliW (Mukherjee et al., 2011; Mukherjee et al., 2016; Altegoer et al., 2016; Oshiro et al., 2019). FliW seems to be the ancestral form of CsrA regulation and is conserved in spirochaetes like *Borrelia burgdorferi* and epsilon-proteobacteria like *Campylobacter jejuni* (Titz et al., 2006; Sze et al., 2011; Dugar et al., 2016; Radomska et al., 2016; Li et al., 2018).

Up to two molecules of FliW can bind to a CsrA dimer but only a single copy is necessary to inhibit RNA binding and does so by a poorly understood mechanism (Altegoer et al., 2016; Oshiro et al., 2019). Genetic analysis indicated that FliW did not bind to the same surface of CsrA that binds RNA and determined that mutation of a remote residue, CsrA N55, abolished both inhibition and FliW interaction (Mukherjee et al., 2016). Similarly, structural analysis indicated that FliW bound to the facing of CsrA opposite of the RNA interaction site, with many contacts near CsrA N55, but how RNA binding was antagonized remained unclear (Altegoer et al., 2016). A model was proposed in which a series of negatively charged residues in an unstructured loop of FliW might electrostatically repel the negatively charged backbone of RNA and prevent CsrA binding (Altegoer et al., 2016). Alternatively, genetic analysis invoked allosteric regulation based on the observation that mutation of one residue buried in the core of the CsrA dimer, CsrA I14, abolished FliW-inhibition but not binding (Mukherjee et al., 2016). Ultimately, both structural and genetic data suggested that FliW binding was necessary but not sufficient to inhibit CsrA, but the mechanism of FliW-mediated inhibition remained unknown.

Here we demonstrate that mutation of the negatively charged series of residues in the FliW loop did not abolish FliW activity and thus the mechanism of inhibition was likely not electrostatic repulsion. We next took an unbiased genetic approach to identify loss-of-function alleles of FliW and found two types of mutations that abolished inhibition of CsrA. The first type of mutation altered residues that were either proximal to, or in contact with, CsrA N55 and disrupted interaction between the two proteins. The second type of mutation altered residues that were proximal to the CsrA dimer core region. A key salt bridge was found to connect residue K123 of FliW and residue D41 of CsrA such that when disrupted, FliW-inhibition was abolished but protein interaction was retained. Moreover, reversing the charged residues at those positions restored inhibition by FliW. The residues required for inhibition formed a linear series of connections in the structure suggesting that binding of FliW to CsrA results in salt-bridge formation, which in turn provokes a conformational change that disrupts either one or both of the RNA binding pockets. We note that the salt bridge appears to be conserved only in the Bacillales, but different core contacts could conceivably substitute in other organisms.

## Results

### Identification of loss-of-function alleles of FliW

Genetic and structural analysis indicated that FliW inhibited CsrA binding to RNA non-competitively by binding a surface of CsrA that does not overlap the RNA-binding site (Fig. 2 and Fig. S4; Mukherjee et al., 2016; Altegoer et al., 2016). One hypothesis for the mechanism of inhibition invoked a loop of the FliW protein with a strong negative charge, which when modeled with RNA bound-CsrA, suggested possible electrostatic interference with the RNA backbone (Fig. 2A; Altegoer et al., 2016). To determine whether the negative loop of FliW was important for the release of flagellin (*hag*) transcript from CsrA, we simultaneously mutated all five negatively charged residues to alanine (*fliW*^*neg5A*^) and tested for the ability to antagonize CsrA phenotypically. Deletion of *fliW* leads to a non-swarming phenotype as uninhibited CsrA restricts flagellin synthesis to levels insufficient for flagellar assembly (Fig. 1A). Ectopic expression of an IPTG-inducible wild type copy of *fliW* restored motility by restoring CsrA antagonism (Fig. 1A). Expression of the *fliW*^*neg5A*^ allele restored motility to a *fliW*-deletion background in a manner indistinguishable from the expression of the wild type (Fig. 1A). We conclude that the negative charges within the loop are not required for FliW to inhibit CsrA. We further conclude that FliW inhibits CsrA through a mechanism other than electrostatic repulsion mediated by the negatively charged loop located near the RNA binding pocket.

**Figure 1.**
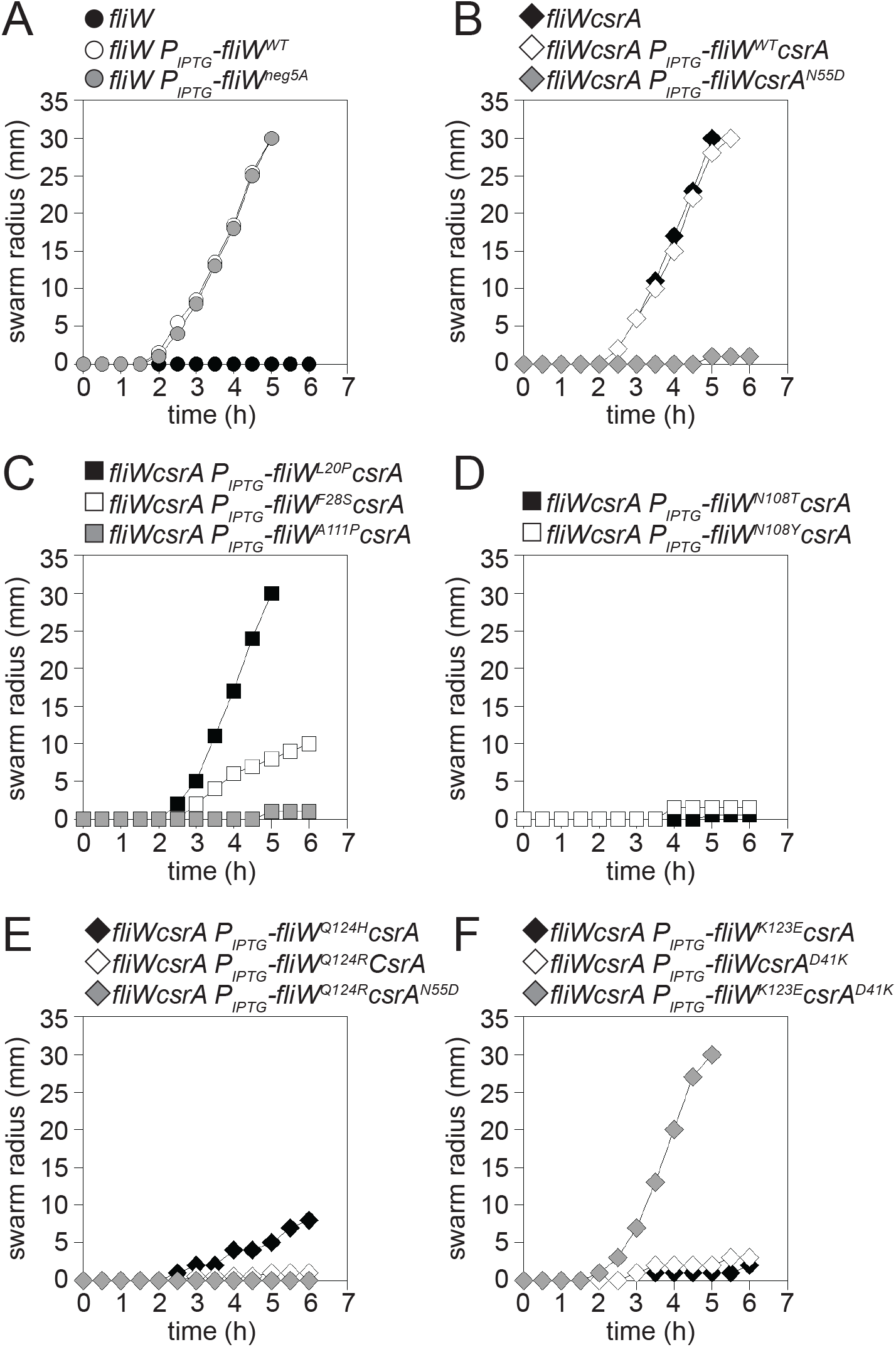
Loss-of-function mutations in *fliW* reduce or abolish swarming motility. A-F) Quantitative swarm expansion assays of artificially induced *fliW* alleles. Each point is the average of three replicates. FliW expression was induced throughout growth and swarming by the addition of 1mM IPTG. The following strains were used to generate the panels: A) *fliW* (DK3237, black circles), *fliW amyE::P*_*IPTG*_-*fliW* (DK2371, white circle), and *fliW amyE::P*_*IPTG*_-*fliW*^*neg5A(E71A,D73A,D75A,E76A,E80A)*^ (DK7342, grey circles). B) *fliWcsrA* (DK2665, black diamonds), *fliWcsrA amyE::P*_*IPTG*_-*fliWcsrA* (DK7567, white diamonds), *fliWcsrA amyE::P*_*IPTG*_-*fliWcsrA*^*N55D*^ (DK8324, grey diamonds). C) *fliWcsrA amyE::P*_*IPTG*_-*fliW*^*L20P*^*csrA* (DK8236, black square), *fliWcsrA amyE::P*_*IPTG*_-*fliW*^*F28S*^*csrA* (DK8244, white squares), and *fliWcsrA amyE::P*_*IPTG*_-*fliW*^*A111P*^*csrA* (DK8237, grey squares). D) *fliWcsrA amyE::P*_*IPTG*_-*fliW*^*N108T*^*csrA* (DK8245, black squares) and *fliWcsrA amyE::P*_*IPTG*_-*fliW*^*N108Y*^*csrA* (DK8326, white squares). E. *fliWcsrA amyE::P*_*IPTG*_-*fliW*^*Q124H*^*csrA* (DK8238, black diamonds), *fliWcsrA amyE::P*_*IPTG*_-*fliW*^*Q124R*^*csrA* (DK8325, white diamonds) and *fliWcsrA amyE::P*_*IPTG*_-*fliW*^*Q124H*^*csrA*^*N55D*^ (DK8345, grey diamonds) F) *fliWcsrA amyE::P*_*IPTG*_-*fliW*^*K123E*^*csrA* (DK7762, black diamonds), *fliWcsrA amyE::P*_*IPTG*_-*fliWcsrA*^*D41K*^ (DK7761, white diamonds), and *fliWcsrA amyE::P*_*IPTG*_-*fliW*^*K123E*^*csrA*^*D41K*^ (DK7717, grey diamonds).

**Figure 2.**
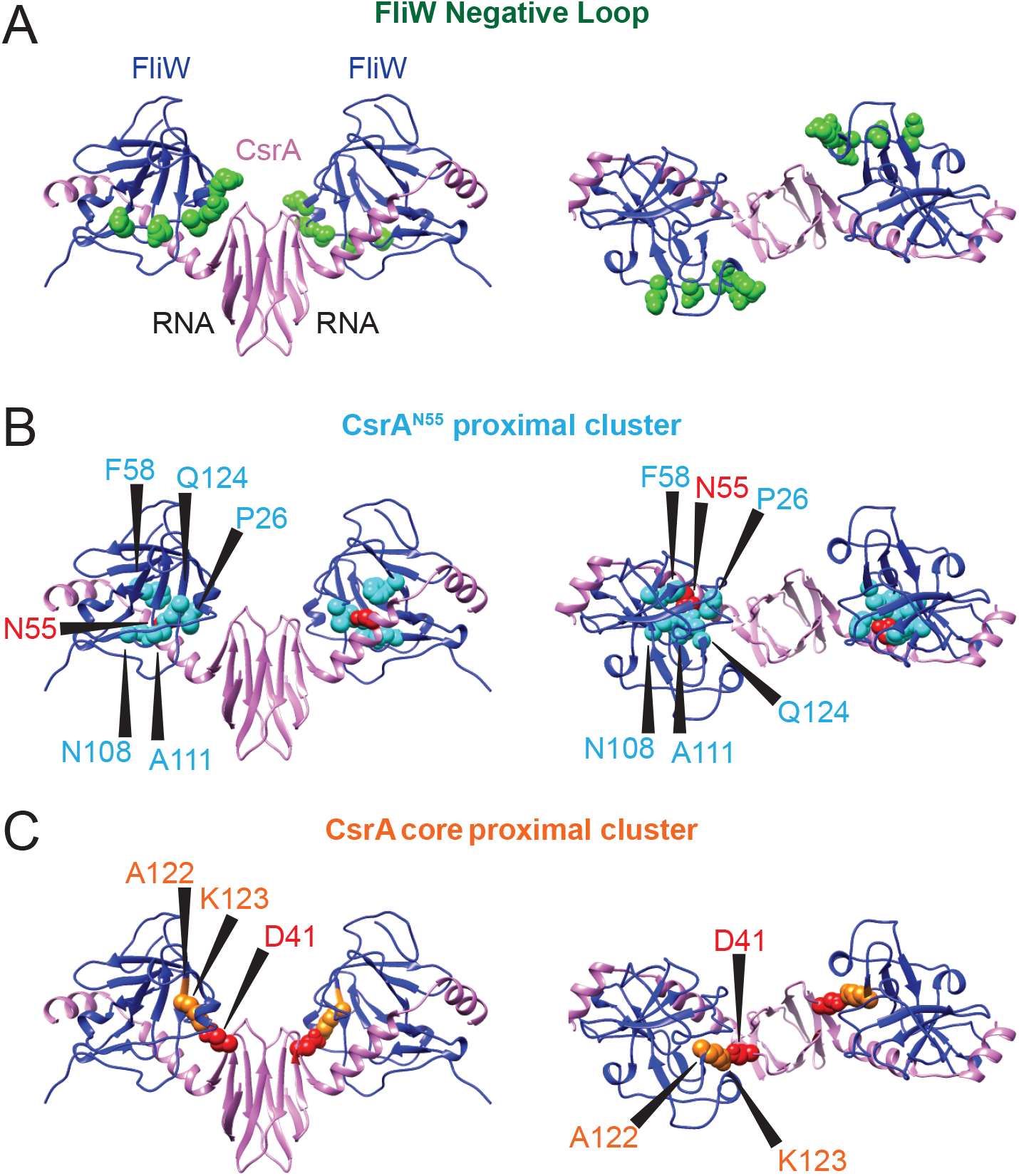
Residues altered by FliW loss-of-function mutations localize to two distinct regions in the FliW-CsrA complex. A-C) Three dimensional structures of the FliW-CsrA complex from *Geobacillus thermodenitrificans* (PDB 5DMB, Altegoer et al., 2016) projected from the “side” (left) and “top” (right) views. The FliW proteins are colored blue and the CsrA dimer is colored lavender. “RNA” indicates the region in which RNA would otherwise be bound. A) Location of the residues constituting the “negative loop”. FliW negatively charged residues within the loop (E71,D73,D75,E76, and E80) are shown in space fill and colored green. B) Location of the residues constituting the “CsrA^N55^ proximal cluster”. FliW residues within the cluster that were mutated (P26, F58, N108, A111, and Q124) are shown in space fill, colored cyan, and indicated by a caret. CsrA residue N55 is shown in space fill and colored red. C) Location of the residues constituting the “CsrA core proximal cluster”. FliW residues within the cluster that were mutated (A122 and K123) are shown in space fill, colored orange, and indicated by a caret. Note that FliW residue 122 of the *G. thermodenitrificans* is a glycine and not an alanine. CsrA residue D41 is shown in space fill and colored red. The location of each residue in FliW which when mutated conferred a loss-of-function mutation can be found individually depicted in Fig. S4.

A forward genetic approach was undertaken to identify loss-of-function mutations in FliW which rendered the protein incapable of inhibiting CsrA. The phenotypic basis for the genetic screen was the aforementioned test for swarming motility (Fig. 1A, Mukherjee et al., 2011). To identify alleles of *fliW* that could not inhibit CsrA, the IPTG-inducible *fliW* construct was mutagenized by error-prone PCR and introduced into the *fliW*-deletion background. The pool was then screened for loss-of-function alleles by gridding individual colonies onto swarm agar and colonies that failed to restore motility upon induction were sought. Over 3,000 colonies were screened and 92 colonies were found to be non-motile suggesting that they encoded loss-of-function alleles in the IPTG-inducible copy of *fliW*.

One way in which an allele of *fliW* would fail to complement the native *fliW* deletion is if the IPTG-inducible copy failed to produce stable FliW protein. Indeed, Western blot analysis indicated that 77 of the 92 candidate isolates failed to produce wild type levels of protein either due to defects in the IPTG-inducible promoter or due to mutations in FliW that reduced protein stability, and all such mutants were discarded from further analysis. The inducible *fliW* construct was sequenced in all 15 of the remaining mutants that produced levels of FliW protein that were either equal to or greater than the amount produced by wild type (Fig. S1 and Fig. S3A). After discarding sibling sequences, two alleles had single point missense mutations (*fliW*^*Q124R*^ and *fliW*^*N108Y*^), and six alleles had double missense mutations (*fliW*^*L20P,A122V*^, *fliW*^*I25T,P26S*^, *fliW*^*F28S,S53P*^, *fliW*^*F58S,Q124H*^, *fliW*^*N108T,M120T,*^ and *fliW*^*A111P,K123E*^).

Previous work indicated that FliW antagonizes CsrA in a 1:1 stoichiometric ratio and the level of FliW to CsrA is maintained through translational coupling (Oshiro et al., 2019). To further validate the loss-of-function alleles of *fliW* while maintaining the appropriate stoichiometric ratio of FliW to CsrA, all alleles were rebuilt as single mutations in a construct that expressed both *fliW* and *csrA* from an IPTG-inducible promoter and integrated at an ectopic site (*amyE::P*_*IPTG*_-*fliWcsrA*), in a *fliWcsrA* double-deletion background (Fig. 1B-F, Fig S2, and Fig. S3B). Both the *fliWcsrA* double-deletion mutant and the double-deletion mutant that was complemented with an IPTG-induced copy of wild type *fliWcsrA* was proficient for motility (Fig. 1B). Motility was lost however, when wild type *fliW* and an allele of *csrA* that cannot be inhibited by FliW (*csrA*^*N55D*^) was co-induced (Fig. 1B). Next, each loss-of-function *fliW* mutation was tested for the ability to maintain motility in the *fliWcsrA* double-deletion background to determine, which if any of the mutations were sufficient to abolish FliW activity (Fig. 1B-F and Fig. S2A-F).

The fourteen reconstructed mutants were separated based on their effect on motility. Six of the mutations (FliW^L20P^, FliW^I25T^, FliW^F28S^ FliW^S53P^, FliW^M120T^, and FliW^Q124H^) had little to no effect on FliW function as indicated by minor changes in motility proficiency and were discarded from further study (Fig. 1C, F and Fig. S2A-F). FliW activity was abolished however in the remaining eight mutants (FliW^P26S^, FliW^F58S^, FliW^N108T^, FliW^N108Y^, FliW^A111P^, FliW^A122V^, FliW^K123E^, FliW^Q124R^) as indicated by the loss of motility (Fig. 1C-F, and Fig. S2A-F). We note that two positions, N108 and Q124, were hit twice with two different amino acid substitutions, and while N108 was allele independent, Q124 was allele-specific such that an arginine substitution abolished activity but a histidine substitution did not (Fig. 1D-E). Ultimately 8 missense alleles that conferred loss-of-function to FliW were retained for further study. We conclude that the ability of FliW to antagonize CsrA can be abrogated or reduced by a variety of single point mutations located throughout the protein.

### Binding of FliW is necessary but not sufficient to inhibit CsrA

To separate the remaining mutations, the location of each mutation was mapped onto the FliW structure within the 3-dimensional FliW-CsrA complex (Altegoer et al., 2016; PDB 5DMB). Most of the *fliW* alleles changed one or more residues (P26, F58, N108, A111P and Q124R), that clustered near residue N55 of CsrA, which was previously identified to be required for inhibition by, and binding of, FliW (Fig. 2B; Mukherjee et al., 2016). Residues A122 and K123 however, were located in a region of FliW that was not near residue N55 of CsrA but rather were proximal to the CsrA core dimerization domain (Fig. 2C). Based on structural analysis, the residues required for inhibition were spatially segregated into two groups, one of which we will refer to as the “CsrA N55 proximal cluster” (Fig. 2B) and the other we will refer to as the “CsrA core proximal cluster” (Fig. 2C). We hypothesized that the two different FliW clusters of residues might be defective in CsrA inhibition for different reasons.

One way in which residues of FliW might be required for CsrA inhibition is if the residues were required for FliW-CsrA interaction. To test for FliW-CsrA interaction *in vivo*, each of the wild type and mutant *fliW* alleles were artificially expressed with *csrA* at an ectopic site of the chromosome in a *fliWcsrAhag* triple mutant background and treated with the crosslinker formaldehyde to trap protein interactions (Fig. 3). Note that FliW also interacts with Hag, and *hag* was deleted in this background as to emphasize FliW-CsrA interaction. The lysates were then resolved on SDS-PAGE and detected by Western blot analysis. In the absence of crosslinker, wild-type FliW and CsrA both ran at their native molecular weights of 16 kDa and 8 kDa respectively (Fig. S5). In the presence of crosslinker however, an additional higher molecular weight band of 24 kDa appeared that contained both FliW and CsrA, trapping a 1:1 FliW:CsrA complex, as previously reported (Fig. 3; Mukherjee et al., 2016). Moreover, the 24kDa band was severely reduced when FliW was expressed with the *csrA*^*N55D*^ allele previously reported to abolish interaction between the two proteins (Fig. 3; Mukherjee et al., 2016). Expression of wild type CsrA with the FliW alleles of the “CsrA N55 proximal cluster”, FliW^P26S^, FliW^F58S^, FliW^N108T^, FliW^N108Y^, FliW^A111P^, and FliW^Q124R^ also reduced or abolished the 24kDa FliW-CsrA complex suggesting that, like CsrA N55 itself, they were required for protein-protein interaction (Fig. 3 and Fig. S5).

**Figure 3.**
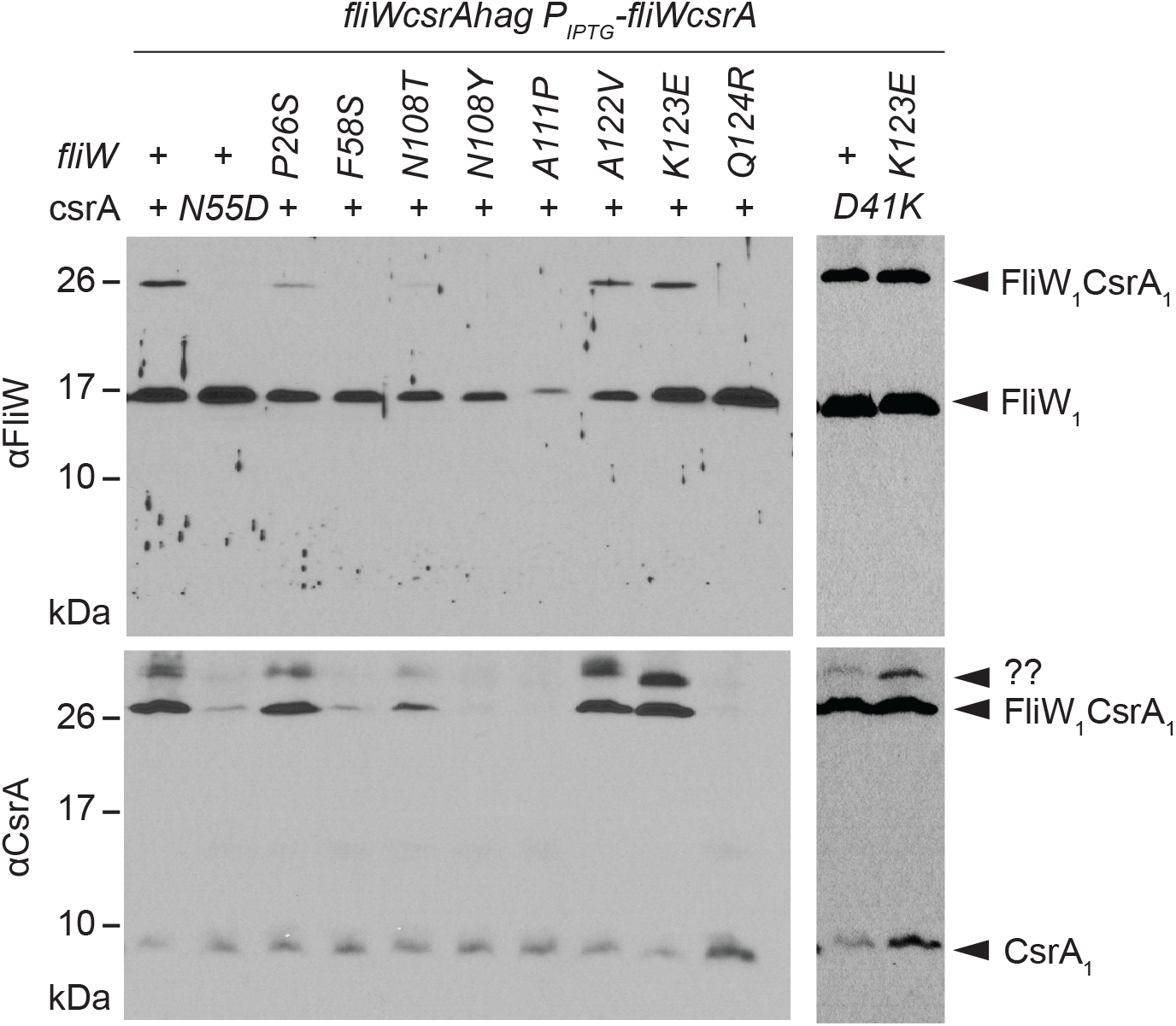
Mutation of FliW “CsrA^N55^ proximal” residues abolish CsrA interaction in vivo but mutation of FliW “CsrA core proximal” residues do not. Whole-cell lysates were grown in the presence of 0.1 mM IPTG, crosslinked with 0.3% formaldehyde, and subjected to Western blot analysis separately using primary antibodies against FliW (Top) and CsrA (Bottom). Samples were prepared from strains containing a triple deletion of *fliW*, *csrA*, and *hag* at their respective native sites (*fliWcsrAhag*). The introduction of the *hag* deletion was to abolish FliW-Hag interaction and reduce the complexity of the interaction species. Translational coupling of *fliW* and *csrA* was maintained by co-expressing both proteins from the same IPTG-inducible promoter integrated at an ectopic site (*amyE::P*_*IPTG*_-*fliWcsrA*). The *csrA* and *fliW* allele is expressed at the top of the panel using either “+” for wild type or the substitution as indicated. The location of CsrA monomer (CsrA1), FliW monomer (FliW1) and FliW-CsrA monomer complex (FliW_1_CsrA_1_) are indicated by carets. CsrA dimer (CsrA2) is not observed, likely because the interaction is too tight to permit formaldehyde crosslinking in this analysis. An additional species (??) is observed in cells expressing FliWK123E but the molecular nature of this species is unknown. The following strains were used to generate lysates for both panels: Top and bottom left) *fliWcsrAhag amyE::P*_*IPTG*_-*fliWcsrA* (DK7974), *fliWcsrAhag amyE::P*_*IPTG*_-*fliWcsrA*^*N55D*^ (DK8303), *fliWcsrAhag amyE::P*_*IPTG*_-*fliW*^*P26S*^*csrA* (DK8282), *fliWcsrAhag amyE::P*_*IPTG*_-*fliW*^*F58S*^*csrA* (DK8284), *fliWcsrAhag amyE::P*_*IPTG*_-*fliW*^*N108T*^*csrA* (DK8304), *fliWcsrAhag amyE::P*_*IPTG*_-*fliW*^*N108Y*^*csrA* (DK8113), *fliWcsrAhag amyE::P*_*IPTG*_-*fliW*^*A111P*^*csrA* (DK8330), *fliWcsrAhag amyE::P*_*IPTG*_-*fliW*^*A122V*^*csrA* (DK8241), *fliWcsrAhag amyE::P*_*IPTG*_-*fliW*^*K123E*^*csrA* (DK8286), and *fliWcsrAhag amyE::P*_*IPTG*_-*fliW*^*Q124R*^*csrA* (DK8108). Top and bottom right) *fliWcsrAhag amyE::P*_*IPTG*_-*fliWcsrA*^*D41K*^ (DK8379) and *fliWcsrAhag amyE::P*_*IPTG*_-*fliW*^*K123E*^*csrA*^*D41K*^ (DK8378). The uncropped image of each Western blot can be seen in Fig. S5.

To further test the interaction between the FliW alleles above and CsrA *in vitro*, a biochemical protein pull-down experiment followed by Western blot analysis was performed (Fig. 4A). GST tagged CsrA (GST-CsrA) was purified and loaded onto glutathione sepharose resin and incubated with either purified untagged wild-type FliW or mutant protein. Two candidate mutants, FliW^N108Y^ and FliW^Q124R^, were chosen because both were altered in residues mutated twice each with two separate substitutions (FliW^N108T/Y^ and FliWQ124^H/R^), and the mutated residues were highly conserved when compared to other FliW homologs (Fig. 5B). After mixing CsrA with FliW, the supernatant was retained, the pellet was washed 4 times and both proteins were eluted by boiling the pelleted resin in SDS-PAGE loading buffer (Fig. 4A). Wild type FliW was retained in the pellet containing GST-CsrA and only accumulated in the supernatant when the amount of FliW was in stoichiometric excess (Fig. 4A). In contrast, both FliW^N108Y^ and FliW^Q124R^ were found abundantly in the supernatant and were absent from the GST-CsrA pellet fraction (Fig. 4A). We conclude that FliW^N108Y^ and FliW^Q124R^ do not interact with CsrA either *in vivo* or *in vitro*, and that residues N108 and Q124 are both required for interaction. We infer that, like the two representative alleles, the cluster of FliW residues proximal to CsrA N55 are all likely important for the interaction between the two proteins.

**Figure 4.**
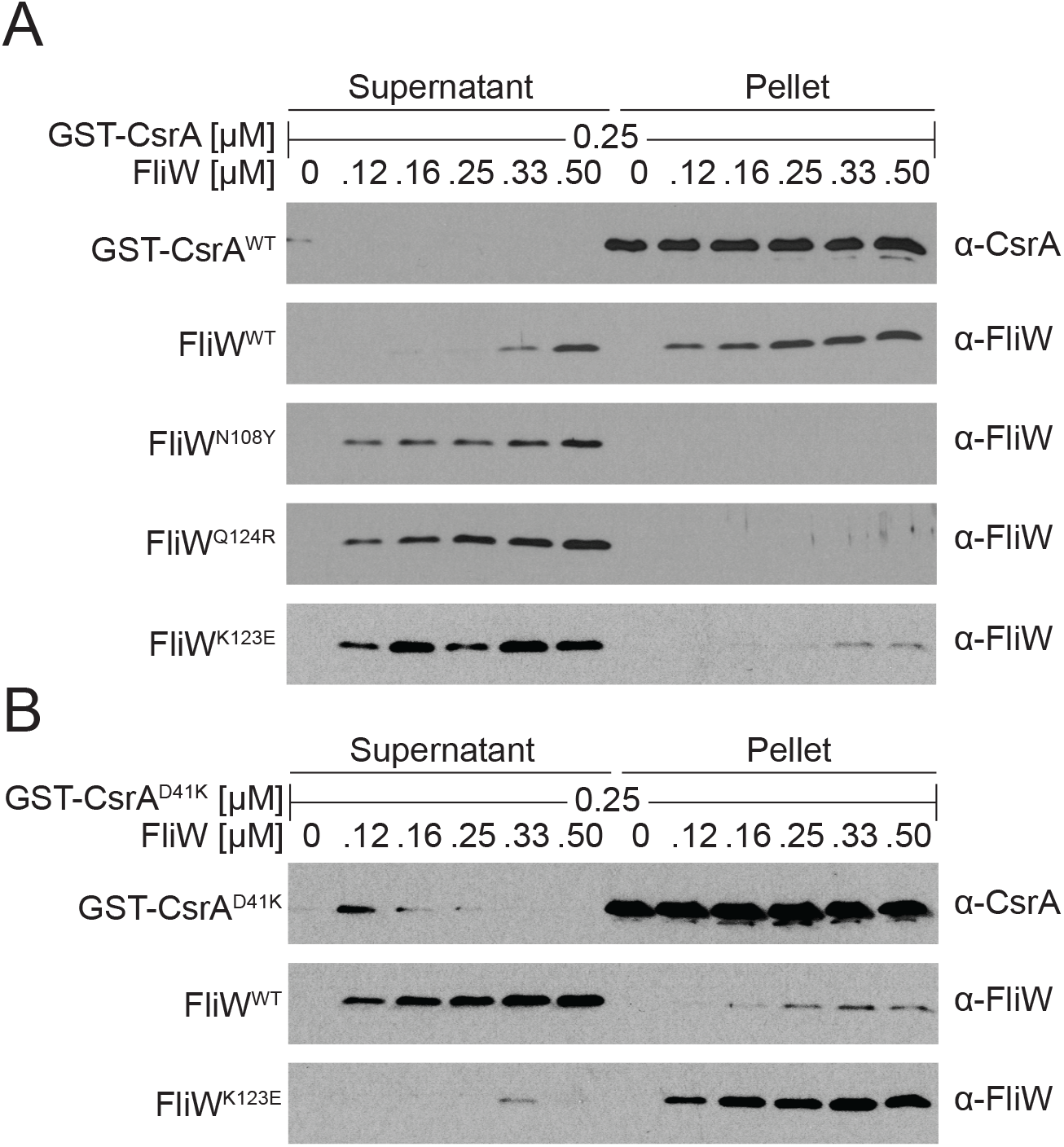
Differential binding of *lof* FliW alleles to CsrA^WT^ and CsrA^D41K^*in vitro*. A-B) A protein pull-down assay using purified GST-CsrA^WT^ or GST-CsrA^D41K^ proteins at the indicated amount was loaded onto a glutathione-sepharose column and incubated with the indicated amounts of purified FliW^WT^, FliW^N108Y^, FliW^Q124R^, or FliW^K123E^ protein in the presence of BSA. Samples were subjected to Western blot analysis and probed with primary antibodies to FliW and CsrA. “Supernatant” indicates the proteins that failed to bind to the beads, and “pellet” indicates the proteins that remained bound to the beads following a series of washes. A) Testing the ability of FliW^WT^ and FliW loss-of-function mutations to interact with GST-CsrA^WT^. B) Testing the ability of FliW and FliW^K123E^ to interact with GST-CsrA^D41K^. Note, that in some instances CsrA can be seen in the supernatant lanes and this is due to resin being accidentally collected while taking the supernatant sample. A representative CsrA blot is shown at the top of A and B. The associated CsrA blot for each FliW allele can be seen in Fig S6.

**Figure 5.**
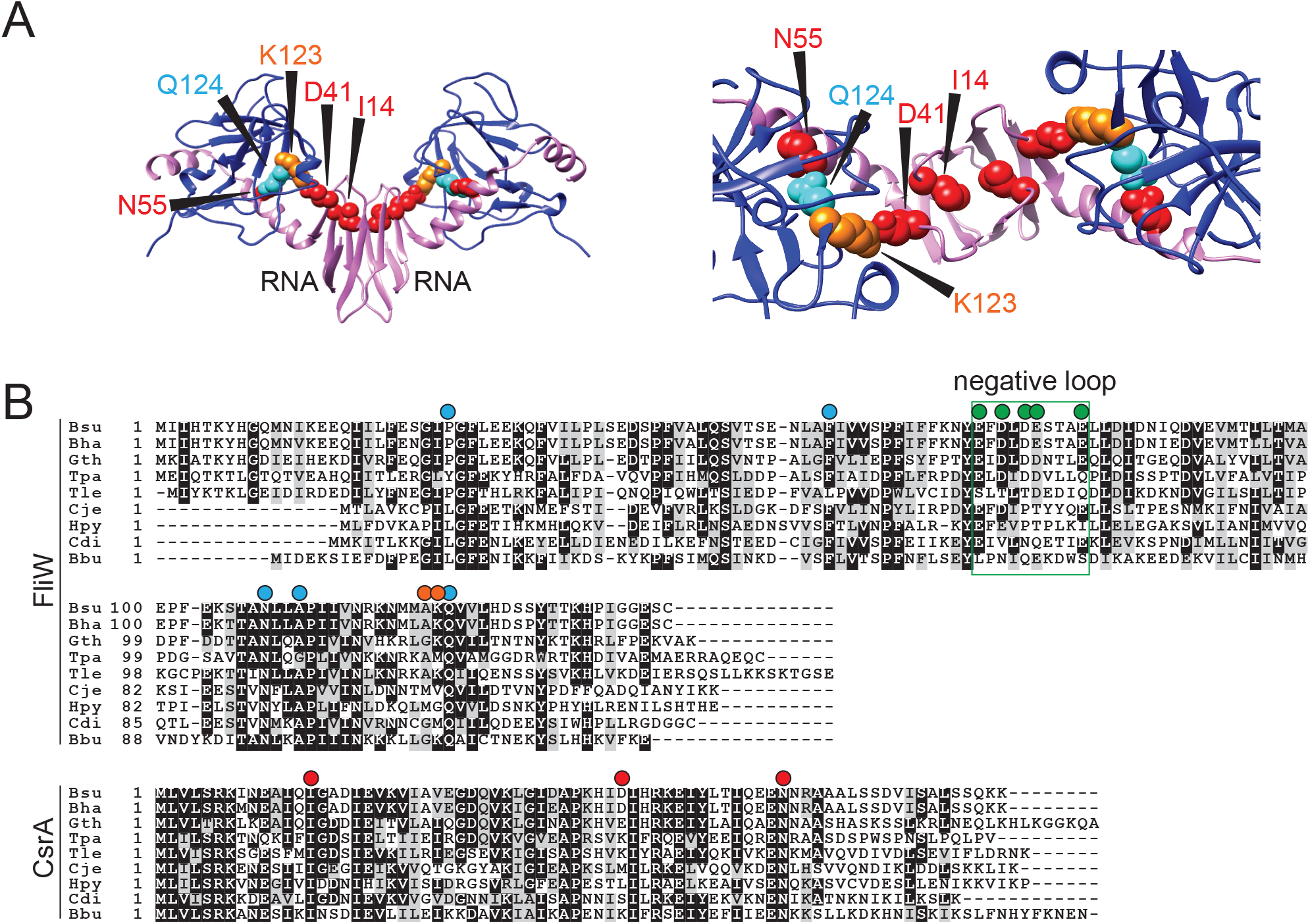
The residues important for CsrA binding and antagonism by FliW form a linear series of connections in space. A) Structural presentation of FliW residues and CsrA residues identified here and reported previously (REF REF REF). The CsrA dimer is colored in lavender and FliW proteins are colored in blue. “RNA” indicates the region in which RNA would otherwise be bound. Side view (left) and a zoomed in view from the top (right) is provided. The FliW residues Q124 (space filled cyan) and K123 (space filled orange) important for CsrA binding and inhibition respectively and CsrA residues N55 and D41 (space filled red) required for binding and inhibition respectively mapped onto the FliW-CsrA complex structure from *G. thermodenitrificans* (PDB 5DMB, REF). CsrA residue I14 (space filled red) is also indicated as mutation at this position to a methionine previously showed that changes to the CsrA core region abolished FliW inhibition but not binding (REF). B) The following organisms were used for a multiple sequence alignment of both FliW (top) and CsrA (bottom): *Bacillus subtilis* (NCIB 3610) “Bsu”, *Bacillus halotolerans* (ATCC 25096) “Bha”, *Geobacillus thermodenitrificans* (NG-80) “Gth”, *Treponema pallidum* (SS14) “Tpa”, *Thermotoga lettingae* (TMO) “Tle”, *Campylobacter jejuni* (ATCC 700819) “Cje”, *Helicobacter pylori* (J99) “Hpy”, *Clostridium difficile* (CD13) “Cdi”, and *Borrelia burgdorferi* (B31) “Bbu”. Positions mutated in *Bacillus subtilis* FliW or CsrA are indicated in above the multiple sequence alignment such that the colored circles that match the colored residues in the structural analysis shown in Fig 2, Fig 5A, and Fig S3. Green circles indicate residues in the FliW “negative loop” (boxed in green). Cyan circles indicate residues in FliW that were mutated near the residue N55 of CsrA “CsrA^N55^ proximal cluster”; orange circles indicate residues that were mutated in FliW near the CsrA core “CsrA core proximal cluster”; red circles indicate residues in CsrA that are either important for FliW binding (N55) or FliW inhibition (I14 and D41).

Structural analysis indicates that residue Q124 of FliW appears to form hydrogen bonds with residue N55 of CsrA suggesting that these two residues form an interaction couple (Fig. 5A). Mutation of either position to a charged residue disrupted interaction, and we wondered whether interaction could be restored if both FliW^Q124R^ and CsrA^N55D^, when expressed in the same cell would restore interaction by the formation of a salt bridge. Simultaneous expression of both mutant alleles failed to support motility in a *fliWcsrA* double-deletion background suggesting that our attempt to artificially replace hydrogen bonding with a salt bridge was unsuccessful to restore productive interaction between the two proteins (Fig. 1E). Expression of FliW^Q124H^ and FliW^Q124K^ with CsrA^N55D^ was similarly insufficient to restore motility (Fig. S2G). Perhaps consistent with the inability to artificially restore interaction, we note that each of these residues are invariant in a sample of FliW-CsrA encoding organisms taken throughout the bacterial phylogeny (Fig. 5B). We conclude that glutamine at position 124 of FliW and asparagine at position 55 of CsrA are critical for the interaction likely because the two residues make direct contact with one another.

### Inhibition of CsrA by FliW depends on the formation of a core-proximal salt bridge

Residues mutated within the FliW “CsrA N55 proximal cluster” reduced or abolished interaction with CsrA, but mutation of residues in the CsrA core proximal cluster preserved interaction with CsrA to levels comparable to that of the wild type when crosslinked *in vivo* (Fig 3). To test for interaction *in vitro*, mutated FliW^K123E^ protein was purified and was mixed with GST-CsrA in a protein pulldown assay and detected by Western blot analysis (Fig. 4A). The FliW^K123E^ protein was predominantly found in the supernatant with little to no FliW visible in the GST-CsrA pellet (Fig. 4A). Thus, mutation of FliW residue K123 to a glutamate did not strictly abolish interaction with CsrA as the interaction could be trapped by a chemical crosslinker. Instead, the residue appeared to be required for stabilizing the complex as the mutant allele was unable to remain in contact with CsrA in a purified *in vitro* system. We conclude that the residues mutated in the FliW “CsrA core proximal cluster” behave differently than residues mutated within the FliW “CsrA N55 proximal cluster” and are likely necessary beyond simple protein interaction.

Structural analysis indicated that the positively charged FliW residue K123 might form a salt bridge with the negatively charged CsrA residue D41 (Fig. 2C and Fig 5A). We note that the FliW^K123E^ allele reversed the charge at that position perhaps resulting in loss of contact with CsrA residue D41 by electrostatic repulsion. Consistent with the idea that the salt bridge is important, motility was inhibited when a form of CsrA was expressed in which residue D41 was changed to an oppositely charged lysine by site-directed mutagenesis (CsrA^D41K^) (Fig. 1F). As with FliW^K123E^, CsrA^D41K^ appeared to maintain FliW-CsrA interaction *in vivo* when trapped by chemical crosslinking (Fig. 3) and poorly retained wild type FliW in the *in vitro* protein pulldown assays (Fig. 4B). Motility was restored however, when both charge-reversed alleles were expressed simultaneously, suggesting that both the salt bridge formation and FliW antagonism had been restored (Fig. 1F). Moreover, combined expression of both mutants also showed interaction in *in vivo* crosslinking Westerns (Fig. 3), and strong interaction between the two alleles was restored in *in vitro* protein pulldowns (Fig. 4B). We conclude that the formation of a salt bridge between FliW and CsrA near the CsrA core region is important for antagonism. We further conclude that this contact plays an allosteric role secondary to the initial interaction event.

## DISCUSSION

CsrA is one of a few proteins in bacteria that bind to transcript RNA and regulate translation (Liu and Romeo, 1997; Dubey et al., 2005., Mercante et al., 2006; Schubert et al., 2007; Mercante et al., 2009). Some bacteria like the gamma-proteobacteria, relieve CsrA inhibition by expressing competitive inhibitor small RNAs with iterations of the CsrA-binding site that act by sequestration (liu et al., 1997; Suzuki et al., 2002; Weilbacher et al., 2003; Kulkarni et al., 2006; Zere et al., 2015). Most other bacteria however, antagonize the RNA-binding activity of CsrA by expressing a protein, FliW (Titz et al., 2006; Sze et al., 2011; Mukherjee et al., 2011; Dugar et al., 2016; Radomska et al., 2016; Li et al., 2018). Genetic and structural analysis supports the notion that FliW functions differently than the sRNA such that CsrA is antagonized non-competitively (Altegoer et al., 2016; Mukherjee et al., 2016). Despite having a high-resolution structure of the inhibited complex, the mechanism of FliW-mediated CsrA-inhibition remains unclear. Here we isolate mutants of FliW defective for antagonism of CsrA and find that the mutants fall into two spatial and functional classes. Combined with previous results, the alleles presented here allow us to expand on a model of FliW interaction that leads to the allosteric inhibition of CsrA.

The foundational observations governing the model of FliW antagonism are as follows. First, FliW and CsrA directly interact, and a highly conserved CsrA residue N55, which when mutated, abolishes both antagonism and interaction simultaneously (Mukherjee et al., 2016; Altegoer et al., 2016). Second, two molecules of FliW may bind to one CsrA dimer but only one molecule of FliW is needed to inhibit interaction with RNA (Altegoer et al., 2016; Oshiro et al., 2019). Third, the structure of the FliW-CsrA complex has been solved, and although it represents the antagonized state, there is little indication of perturbation to, or occlusion of, the RNA binding pocket (Altegoer et al., 2016). Importantly, interaction is necessary but not sufficient for antagonism, as a mutation of a particular residue within the CsrA core (I14) to a bulkier methionine was found to abolish antagonism while retaining the ability to bind FliW (Mukherjee et al., 2016). How alteration of CsrA residue I14 abolishes antagonism is unclear but seems to suggest that the conformation of CsrA is important. Thus, FliW binds to a surface of CsrA that is remote from the active site, but antagonism requires an additional step.

To add to these observations, we took an unbiased approach and identified the residues of FliW that are required for inhibition. One group of required FliW residues were proximal to the regulatory CsrA residue N55 (Fig. 2B). Similar to mutation of CsrA residue N55 itself, mutation of the proximal FliW residues abolished both antagonism and interaction between the two proteins as observed in both *in vivo* and *in vitro* assays (Fig. 3 and Fig. 4A). FliW residue Q124 in particular appears to make direct contact with CsrA residue N55 in the FliW-CsrA complex, and we note that both residues appear to be invariant amongst different homologs in genomes where both proteins are encoded (Fig. 5; Altegoer et al., 2016). Moreover, mutation of these residues was allele-independent and substitutions that could potentially restore interaction between these two residues were not tolerated (Fig 1E and Fig. S2G). We suspect that the intolerance is likely due to the wide variety of conserved FliW residues required within the “CsrA N55 proximal” cluster, each of which seem to contribute to contact and may therefore exclude non-native combinations of position 124 of FliW and position 55 of CsrA (Fig. 2B and Fig. 5B). Whatever the case, we conclude the FliW residues that make up “CsrA N55 proximal cluster” are required for antagonism because they mediate the initial interaction between the two proteins.

Another group of required FliW residues were proximal to the CsrA core region (Fig. 2C). Mutation of the core proximal residues abolished antagonism but behaved more like the CsrA^I14M^ allele in that they maintained the ability to bind FliW (Mukherjee et al., 2016). One residue in particular, FliW K123, appeared to form a salt bridge connection with CsrA residue D41, and mutation of either residue to the opposite charge maintained complex formation between the two proteins *in vivo* when trapped by chemical crosslinking (Fig. 3 and Fig. 5A). We note that each charge reversed allele decreased complex stability *in vitro*, but the reduced stability could have been due to the introduction of non-native electrostatic repulsion (Fig. 4). Charge reversal on both proteins simultaneously however, restored FliW-mediated inhibition suggesting that the precise residues were less important than the restoration of the quaternary structure interaction at that position (Fig. 1F and Fig. 4B). Consistent with allele flexibility, we note that the residues at FliW position 123 and CsrA position 41 vary in genomes that express both proteins and indeed, the salt bridge seems to be particular to the Bacillales (Fig. 5B). The interaction with the CsrA core domain seems to be what is important and this could be accomplished by different residues at perhaps slightly different locations in other organisms.

The combined genetic data indicates that the required residues of *B. subtilis* FliW connect two distinct regions of CsrA as a linear chain (Fig. 5A). We suggest that initial binding is mediated by direct interaction between CsrA residue N55 and FliW residue Q124. The adjacent residue of FliW K123 then forms a salt bridge with CsrA D41 resulting in allosteric inhibition of RNA binding activity. The mechanism for allosteric regulation during the second step of FliW antagonism is still unclear. The allosteric salt bridge could change the FliW structure, perhaps by moving the negative loop into the RNA binding pocket of CsrA, and while steric inhibition is possible, here we show that the electrostatic repulsion is dispensible. We note that a change in FliW conformation upon binding is unlikely however, as the structure of FliW on its own is the same as when it is bound to CsrA (Altegoer et al., 2016). Instead we favor a model in which CsrA changes conformation because the linear series of amino acids that mediate the inhibition are in line with the CsrA core residue I14 (Fig. 5A). We note that the CsrA^I14M^ allele phenocopies disruption of the salt bridge and we speculate that changing to a bulkier methionine prevents deformation of, and allosteric conformational change through, the core domain following FliW binding (Mukherjee et al., 2016). In toto, our results provide further evidence that FliW antagonism occurs in two steps with the second step playing an allosteric role.

## MATERIALS AND METHODS

### Strains and growth conditions

*B. subtilis* strains were grown in lysogeny broth (LB) (10 g tryptone, 5 g yeast extract, 5 g NaCl per L) broth or on LB plates fortified with 1.5% Bacto agar at 37°C. When appropriate, antibiotics were included at the following concentrations: 10 μg/ml tetracycline (tet), 100 μg/ml spectinomycin (spec), 5 μg/ml chloramphenicol (cm), 5 μg/ml kanamycin (kan), and 1 μg/ml erythromycin plus 25 μg/ml lincomycin (*mls*). Isopropyl β-D-thiogalactopyranoside (IPTG, Sigma) was added to the medium at the indicated concentration when appropriate.

For quantitative Swarm expansion assay, strains were grown to mid-log phase (OD_600_ 0.3 – 1.0) and concentrated to 10 OD_600_ in PBS pH 7.4 (137 mM NaCl, 2.7 mM KCl, 10 mM Na_2_HPO_4_, and 2 mM KH_2_PO_4_) containing 0.5% India ink (Higgins). LB plates containing 0.7% Bacto agar with or without varying concentrations of IPTG were dried for 10 minutes in a laminar flow hood, centrally inoculated with 10 μl of the cell suspension, dried for another 10 minutes, and incubated at 37°C in a humid chamber. Swarm radius was measured along the same axis every 30 min.

### Strain construction

All constructs were either introduced into a 3610-derived natural competent strain DK1042 (Konkol et al., 2013) or first introduced by natural competence into a domesticated strain PY79 or a 3610-derived competent strain cured of the pBS32 plasmid DS2569 (Konkol et al., 2013) then transferred to the 3610 background using SPP1-mediated generalized phage transduction (Yasbin and Young., 1974). Briefly, SPP1-mediated transduction were performed by generating a lysate on *B. subtilis* grown in TY (1% Tryptone, 0.5% yeast extract, 0.5% NaCl, 10 mM MgSO_4_, and 1 mM MnSO4). Recipient strains were grown to stationary phase in TY, 1 ml diluted into 9 ml TY, and 25 μl (for streptomycin) lysates were added, followed by incubation at room temperature for 30 min, and then selection on the respective antibiotic at 37°C overnight. For transductions in which spectinomycin-resistance was selected for, 10 mM sodium citrate was added to the selection plate. All strains used in this study are listed in Table 1. All primers used in this study are listed in supplemental table S1. All plasmids used in this study are listed in supplemental table S2.

**Table 1:** Strains

| Strain | Genotype |
| --- | --- |
| 3610 | Wild type |
| DK1042 | *comI*<sup>Q12L</sup> (Konkol et al., 2013) |
| DK1483 | *comI*<sup>Q12L</sup> Δ*fliWcsrAhag* (Mukherjee et al., 2016) |
| DK2371 | Δ*fliW amyE::P<sub>hyspank</sub>-fliW spec* (Mukherjee et al., 2011) |
| DK2665 | *comI*<sup>Q12L</sup> Δ*fliWcsrA* (Oshiro et al., 2019) |
| DK3237 | *comI*<sup>Q12L</sup> Δ*fliW* |
| DK4205 | *comI*<sup>Q12L</sup> *csrA*<sup>N55D</sup> (Oshiro et al., 2019) |
| DK6425 | *comI*<sup>Q12L</sup> Δ*fliW amyE::P<sub>hyspank</sub>-fliW*<sup>N108Y</sup> *spec* |
| DK6426 | *comI*<sup>Q12L</sup> Δ*fliW amyE::P<sub>hyspank</sub>-fliW*<sup>F28S,S53P</sup> *spec* |
| DK6428 | *comI*<sup>Q12L</sup> Δ*fliW amyE::P<sub>hyspank</sub>-fliW*<sup>N108T,M120T</sup> *spec* |
| DK7254 | *amyE::P<sub>hyspank</sub>-fliW*<sup>E71A,D73A,D75A,E76A,E80A</sup> *spec* |
| DK7342 | *comI*<sup>Q12L</sup> Δ*fliW amyE::P<sub>hyspank</sub>-fliW*<sup>E71A,D73A,D75A,E76A,E80A</sup> *spec* |
| DK7386 | *comI*<sup>Q12L</sup> Δ*fliW amyE::P<sub>hyspank</sub>-fliW*<sup>F58S,Q124H</sup> *spec* |
| DK7387 | *comI*<sup>Q12L</sup> Δ*fliW amyE::P<sub>hyspank</sub>-fliW*<sup>L20P,A122V</sup> *spec* |
| DK7389 | *comI*<sup>Q12L</sup> Δ*fliW amyE::P<sub>hyspank</sub>-fliW*<sup>Q124R</sup> *spec* |
| DK7567 | *comI*<sup>Q12L</sup> Δ*fliWcsrA amyE::P<sub>hyspank</sub>-fliWcsrA spec* |
| DK7714 | *amyE::P<sub>hyspank</sub>-fliW*<sup>K123E</sup> *csrA*<sup>D41K</sup> *spec* |
| DK7717 | *comI*<sup>Q12L</sup> Δ*fliWcsrA amyE::P<sub>hyspank</sub>-fliW*<sup>K123E</sup> *csrA*<sup>D41K</sup> *spec* |
| DK7741 | *comI*<sup>Q12L</sup> Δ*fliW amyE::P<sub>hyspank</sub>-fliW*<sup>A111P,K123E</sup> *spec* |
| DK7742 | *comI*<sup>Q12L</sup> Δ*fliW amyE::P<sub>hyspank</sub>-fliW*<sup>I25T,P26S</sup> *spec* |
| DK7747 | *amyE::P<sub>hyspank</sub>-fliWcsrA*<sup>D41K</sup> *spec* |
| DK7761 | *comI*<sup>Q12L</sup> Δ*fliWcsrA amyE::P<sub>hyspank</sub>-fliWcsrA*<sup>D41K</sup> *spec* |
| DK7762 | *comI*<sup>Q12L</sup> Δ*fliWcsrA amyE::P<sub>hyspank</sub>-fliW*<sup>K123E</sup> *csrA spec* |
| DK7974 | *comI*<sup>Q12L</sup> Δ*fliWcsrAhag amyE::P<sub>hyspank</sub>-fliWcsrA spec* |
| DK7997 | *amyE::P<sub>hyspank</sub>-fliWcsrA*<sup>N55D</sup> *spec* |
| DK8085 | *amyE::P<sub>hyspank</sub>-fliW*<sup>Q124R</sup> *csrA spec* |
| DK8090 | *amyE::P<sub>hyspank</sub>-fliW*<sup>N108Y</sup> *csrA spec* |
| DK8108 | *comI*<sup>Q12L</sup> Δ*fliWcsrAhag amyE::P<sub>hyspank</sub>-fliW*<sup>Q124R</sup> *csrA spec* |
| DK8113 | *comI*<sup>Q12L</sup> Δ*fliWcsrAhag amyE::P<sub>hyspank</sub>-fliW*<sup>N108Y</sup> *csrA spec* |
| DK8206 | *amyE::P<sub>hyspank</sub>-fliW*<sup>L20P</sup> *csrA spec* |
| DK8207 | *amyE::P<sub>hyspank</sub>-fliW*<sup>A111P</sup> *csrA spec* |
| DK8208 | *amyE::P<sub>hyspank</sub>-fliW*<sup>Q124H</sup> *csrA spec* |
| DK8209 | *amyE::P<sub>hyspank</sub>-fliW*<sup>I25T</sup> *csrA spec* |
| DK8210 | *amyE::P<sub>hyspank</sub>-fliW*<sup>P26S</sup> *csrA spec* |
| DK8211 | *amyE::P<sub>hyspank</sub>-fliW*<sup>A122V</sup> *csrA spec* |
| DK8212 | *amyE::P<sub>hyspank</sub>-fliW*<sup>F58S</sup> *csrA spec* |
| DK8213 | *amyE::P<sub>hyspank</sub>-fliW*<sup>S53P</sup> *csrA spec* |
| DK8214 | *amyE::P<sub>hyspank</sub>-fliW*<sup>F28S</sup> *csrA spec* |
| DK8215 | *amyE::P<sub>hyspank</sub>-fliW*<sup>N108T</sup> *csrA spec* |
| DK8216 | *amyE::P<sub>hyspank</sub>-fliW*<sup>M120T</sup> *csrA spec* |
| DK8236 | *comI*<sup>Q12L</sup> Δ*fliWcsrA amyE::P<sub>hyspank</sub>-fliW*<sup>L20P</sup> *csrA spec* |
| DK8237 | *comI*<sup>Q12L</sup> Δ*fliWcsrA amyE::P<sub>hyspank</sub>-fliW*<sup>A111P</sup> *csrA spec* |
| DK8238 | *comI*<sup>Q12L</sup> Δ*fliWcsrA amyE::P<sub>hyspank</sub>-fliW*<sup>Q124H</sup> *csrA spec* |
| DK8239 | *comI*<sup>Q12L</sup> Δ*fliWcsrA amyE::P<sub>hyspank</sub>-fliW*<sup>I25T</sup> *csrA spec* |
| DK8240 | *comI*<sup>Q12L</sup> Δ*fliWcsrA amyE::P<sub>hyspank</sub>-fliW*<sup>P26S</sup> *csrA spec* |
| DK8241 | *comI*<sup>Q12L</sup> Δ*fliWcsrA amyE::P<sub>hyspank</sub>-fliW*<sup>A122V</sup> *csrA spec* |
| DK8242 | *comI*<sup>Q12L</sup> Δ*fliWcsrA amyE::P<sub>hyspank</sub>-fliW*<sup>F58S</sup> *csrA spec* |
| DK8243 | *comI*<sup>Q12L</sup> Δ*fliWcsrA amyE::P<sub>hyspank</sub>-fliW*<sup>S53P</sup> *csrA spec* |
| DK8244 | *comI*<sup>Q12L</sup> Δ*fliWcsrA amyE::P<sub>hyspank</sub>-fliW*<sup>F28S</sup> *csrA spec* |
| DK8245 | *comI*<sup>Q12L</sup> Δ*fliWcsrA amyE::P<sub>hyspank</sub>-fliW*<sup>N108T</sup> *csrA spec* |
| DK8246 | *comI*<sup>Q12L</sup> Δ*fliWcsrA amyE::P<sub>hyspank</sub>-fliW*<sup>M120T</sup> *csrA spec* |
| DK8282 | *comI*<sup>Q12L</sup> Δ*fliWcsrAhag amyE::P<sub>hyspank</sub>-fliW*<sup>P26S</sup> *csrA spec* |
| DK8283 | *comI*<sup>Q12L</sup> Δ*fliWcsrAhag amyE::P<sub>hyspank</sub>-fliW<sup>A122V</sup>csrA spec* |
| DK8284 | *comI*<sup>Q12L</sup> Δ*fliWcsrAhag amyE::P<sub>hyspank</sub>-fliW<sup>F58S</sup>csrA spec* |
| DK8286 | *comI*<sup>Q12L</sup> Δ*fliWcsrAhag amyE::P<sub>hyspank</sub>-fliW<sup>K123E</sup>csrA spec* |
| DK8303 | *comI*<sup>Q12L</sup> Δ*fliWcsrAhag amyE::P<sub>hyspank</sub>-fliWcsrA<sup>N55D</sup> spec* |
| DK8304 | *comI*<sup>Q12L</sup> Δ*fliWcsrAhag amyE::P<sub>hyspank</sub>-fliW<sup>N108T</sup>csrA spec* |
| DK8324 | *comI*<sup>Q12L</sup> Δ*fliWcsrA amyE::P<sub>hyspank</sub>-fliWcsrA<sup>N55D</sup> spec* |
| DK8325 | *comI*<sup>Q12L</sup> Δ*fliWcsrA amyE::P<sub>hyspank</sub>-fliW<sup>Q124R</sup>csrA spec* |
| DK8326 | *comI*<sup>Q12L</sup> Δ*fliWcsrA amyE::P<sub>hyspank</sub>-fliW<sup>N108Y</sup>csrA spec* |
| DK8330 | *comI*<sup>Q12L</sup> Δ*fliWcsrAhag amyE::P<sub>hyspank</sub>-fliW<sup>A111P</sup>csrA spec* |
| DK8335 | *amyE::P<sub>hyspank</sub>-fliW<sup>Q124R</sup>csrA<sup>N55D</sup> spec* |
| DK8336 | *amyE::P<sub>hyspank</sub>-fliW<sup>Q124K</sup>csrA spec* |
| DK8337 | *amyE::P<sub>hyspank</sub>-fliW<sup>Q124K</sup>csrA<sup>N55D</sup> spec* |
| DK8345 | *comI*<sup>Q12L</sup> Δ*fliWcsrA amyE::P<sub>hyspank</sub>-fliW<sup>Q124R</sup>csrA<sup>N55D</sup> spec* |
| DK8346 | *comI*<sup>Q12L</sup> Δ*fliWcsrA amyE::P<sub>hyspank</sub>-fliW<sup>Q124K</sup>csrA spec* |
| DK8347 | *comI*<sup>Q12L</sup> Δ*fliWcsrA amyE::P<sub>hyspank</sub>-fliW<sup>Q124K</sup>csrA<sup>N55D</sup> spec* |
| DK8373 | *amyE::P<sub>hyspank</sub>-fliW<sup>Q124H</sup>csrA<sup>N55D</sup> spec* |
| DK8378 | *comI*<sup>Q12L</sup> Δ*fliWcsrAhag amyE::P<sub>hyspank</sub>-fliW<sup>K123E</sup>csrA<sup>D41K</sup> spec* |
| DK8379 | *comI*<sup>Q12L</sup> Δ*fliWcsrAhag amyE::P<sub>hyspank</sub>-fliWcsrA<sup>D41K</sup> spec* |
| DK8380 | *comI*<sup>Q12L</sup> Δ*fliWcsrA amyE::P<sub>hyspank</sub>-fliW<sup>Q124H</sup>csrA<sup>N55D</sup> spec* |
| DS2569 | Δ*pBS32* (Konkol et al., 2013) |
| PY79 | Wild type |

#### In-frame deletions

To generate the *∆fliW* in frame marker-less deletion construct in the DK1042 background, plasmid pJP87 (Mukherjee et al., 2011) was introduced and plasmid integration was selected for by *mls* resistance at 37°C. Plasmid pJP87 encodes a temperature sensitive origin that is active at room temperature but not at 37°C. To evict the plasmid, the strain was subsequently incubated at a room temperature overnight without antibiotic selection. Cells were then serially diluted and plated on LB agar at 37°C. Individual colonies were patched on LB plates and LB plates containing *mls* to identify *mls* sensitive colonies that had evicted the plasmid. Deletion of *fliW* was verified by PCR using primers 1541/1544.

#### Generation of *fliW* mutant pool

To generate a pool of *fliW* mutants, primer pair 861/862 was used to amplify the *fliW* reading frame, using DK2371 chromosomal DNA as a template and Expand polymerase with Expand buffer 2 (Roche)/ The mutant PCR library was first transformed into PY79 by natural competence, and phage lysates were generated from the resulting transformants. The phage lysates were used to transduce strains of *B. subtilis* for screening of loss-of-function alleles.

#### *P*_*hyspank*_-*fliW*^*neg5A*^ construct

Site direct mutations were introduced into the *P*_*hyspank*_-*fliW spec* construct in a stepwise fashion using DK2371 genomic DNA, amplified using primers pairs in 861/6805 862/6806 for *fliW*^*E71A,D73A*^, 861/5871 862/5872 for *fliW*^*D75A,E76A*^, and 861/6707 862/6708 for *fliW*^*E80A*^. The two fragments were assembled together by isothermal “Gibson” assembly (Gibson et al., 2009), introduced into laboratory strain PY79 by natural competence. Mutations were verified prior to introduction of the next set of mutations by amplifying the appropriate gDNA with primer pair 6275/689 and sequenced using primers 1541 and 1542. Once all mutations were confirmed, the construct was introduced into the appropriate strain backgrounds by SPP1-mediated transductions (Yasbin and Young., 1974).

#### *P*_*hyspank*_-*fliWcsrA* mutation constructs

Site directed mutations to *the P*_*hyspank*_-*fliWcsrA spec* construct were generated using DK7567 genomic DNA, amplified using primer pairs 861/7098 862/7099 for *fliW*^*L20P*^, 861/7102 862/7103 for *fliW*^*A111P*^, 861/7104 862/7105 for *fliW*^*K123E*^, 861/7110 862/7111 for *fliW*^*Q124H*^, 861/7112 862/7113 for *fliW*^*I25T*^, 861/7114 862/7115 for *fliW*^*P26S*^, 861/7116 862/7117 for *fliW*^*A122V*^, 861/7118 862/7119 for *csrA*^*D41K*^, 861/7163 862/7164 for *fliW*^*F58S*^, 861/7165 862/7166 for *fliW*^*S53P*^, 861/7256 862/7257 for *fliW*^*Q124R*^, 861/7258 862/7259 for *fliW*^*F28S*^, 861/7260 862/7261 for *fliW*^*N108T*^, 861/7262 862/7263 for *fliW*^*M120T*^, 861/7266 862/7267 for *fliW*^*N108Y*^, and 861/7311 862/7312 *fliW*^*Q124K*^. The two fragments were assembled together by isothermal “Gibson” assembly (Gibson et al., 2009), introduced into laboratory strain PY79 by natural competence, and further introduced into the appropriate strain backgrounds by SPP1-mediated transductions (Yasbin and Young., 1974). Mutations were verified by amplifying the appropriate gDNA with primer pair 6275/689, and sequenced using primers 1541 and 1544. Construct *P*_*hyspank*_-*fliW*^*K123E*^*csrA*^*D41K*^ was generated using DK7762 chromosomal DNA and amplified using primers pairs 861/7118 and 862/7119, construct *P*_*hyspank*_-*fliW*^*Q124R*^*csrA*^*N55D*^ was generated using DK8324 chromosomal DNA and amplified using primers pairs 861/7256 and 862/7257, construct *P*_*hyspank*_-*fliW*^*Q124H*^*csrA*^*N55D*^ was generated using DK8324 chromosomal DNA and amplified using primers pairs 861/7110 and 862/7111, and construct *P*_*hyspank*_-*fliW*^*Q124K*^*csrA*^*N55D*^ was generated using DK8324 chromosomal DNA and amplified using primers pairs 861/7311 and 862/7312. The previous four constructs were assembled and introduced into the appropriate strain backgrounds as above.

#### *P*_*hyspank*_-*fliWcsrA*^*N55D*^ construct

To generate Physpank-fliWcsrAN55D, a fragment containing fliWcsrA^N55D^ was amplified by using DK4025 as a template and primer pair 1541/1544 and was digested with NheI/SphI. The fragment was ligated into the NheI and SphI sites of pDR11 containing an ampicillin and spectinomycin resistance cassette, to generate pRO102. pRO102 was introduced into the laboratory strain PY79 by natural transformation, then further introduced into the appropriate strain background using SPP1-mediated transduction (Yasbin and Young., 1974).

#### GST-CsrA^D41K^ expression vector

To generate a translational fusion of CsrA^D41K^ to the GST tag, a fragment containing *csrA*^D41K^ was amplified by using DK7761 as a template and primer pair 2140/2141 and was digested with BamHI/EcoRI. The fragment was ligated into the BamHI and EcoR1 sites of pGEX-2TK containing an ampicillin resistance cassette, to create pRO114.

#### His-SUMO-FliW^mutant^ expression vectors

To generate a translational fusion of FliWN108Y, FliWQ124R, and FliWK123E to the His-SUMO tag, a fragment containing the aforementioned mutations was amplified by using DK8326 (*fliW*^*N108Y*^), DK8325 (*fliW*^*Q124R*^), and DK7762 (*fliW*^*K123E*^) as a template and primer pair 2230/7194 and was digested with SapI/XhoI. The fragment was ligated into the SapI and XhoI sites of pTB146 containing an ampicillin resistance cassette, to create pRO106, pRO107, and pRO115.

### GST-CsrA^WT^ and GST-CsrA^D41K^ protein purification

The GST-CsrA^WT^ and GST-CsrA^D41K^ protein expression vectors pSM6 and pRO114 respectively, were transformed into Rosetta gami *E. coli*, grown to ~0.7 OD600 in 500 ml of Luria-Bertani Broth, induced with 1 mM IPTG and grown for 3 h at 37°C. Cells were pelleted and resuspended in Lysis Buffer (25 mM Tris-HCl (pH 8.0), 1 mM DTT, 1 mM EDTA, 150 mM NaCl, 0.5 mM PMSF), and frozen at −80°C for overnight. The frozen cell pellet was thawed and lysed by sonication. Lysed cells were ultracentrifuged at 14000 rpm for 30 min at 4°C. Cleared supernatant was combined with Glutathione-Sepharose (GE Healthcare) and incubated for 3 h at 4°C. The bead/lysate mixture was poured onto a 1 cm separation column (Bio-Rad), the resin was allowed to pack and was washed with Wash Buffer (25 mM Tris-HCl [pH 8.0], 1 mM DTT, 1 mM EDTA, 250 mM NaCl, 10% glycerol, 0.5 mM PMSF). GST-CsrA bound to the resin was then eluted using GST-elution buffer (25 mM Tris-HCl [pH 8.5], 20 mM Glutathione, 1 mM DTT, 1 mM EDTA, 250 mM NaCl, 10% glycerol, 0.5 mM PMSF). Elutions were separated by SDS-PAGE and Coomassie stained to verify purification of the GST-CsrA fusion and the appropriate fractions were pooled and concentrated to ~2mL. Final purification of GST-CsrA protein was conducted via size exclusion chromatography on a Superdex 75 16/60 (GE Healthcare) column using CsrA gel filtration buffer (20 mM Tris-HCl pH 8.0, 200 mM NaCl, 10% Glycerol, 1mM EDTA pH 8.0) and fractions were concentrated and stored at −80C. Concentration of GST-CsrA and GST-CsrA^D41K^ was determined by Bradford assay (Biorad).

### His_6_-SUMO-FliW^WT^ and -FliW^mutant^ Protein Purification

The His_6_-SUMO-FliW^WT^, FliW^N108Y^, FliW^K123E^, and FliW^Q124R^ protein expression vectors pSM12, pRO115, pRO107, and pRO106 were transformed into Rosetta gami *E. coli,* respectively. Cells were grown to **∼**0.7 OD600 in 500 mL of terrific broth, induced with 1 mM IPTG and grown overnight at 16 °C. Cells were pelleted and resuspended in CsrA lysis buffer (100 mM Tris-HCl pH 8.0 and 400 mM NaCl), treated with lysozyme and lysed by sonication. Lysed cells were centrifuged at 14,000 × g for 30 min. Cleared supernatant was combined with Ni-NTA resin (Novagen) and incubated for 1 h at 4 °C. The bead/lysate mixture was poured onto a 1-cm separation column (Bio-Rad), the resin was allowed to pack, and was washed with CsrA Wash Buffer (50 mM Tris-HCl pH 8.0, 200 mM NaCl, and 10% Glycerol). His_6_-SUMO-FliW^WT/N108Y/K123E/Q124R^ bound to the resin was then eluted using a stepwise elution of CsrA Wash Buffer with 5, 15, and 500 mM imidazole. Eluted proteins were separated by SDS/PAGE and Coomassie stained to verify purification of the His_6_-SUMO-FliW^WT/N108Y/K123E/Q124R^ and appropriate fractions were pooled and concentrated to ~2mL. His_6_-SUMO-FliW^WT/N108Y/K123E/Q124R^ protein was further cleaned by size exclusion chromatography on a Superdex 75 16/60 (GE Healthcare) column using CsrA gel filtration buffer (20 mM Tris-HCl pH 8.0, 200 mM NaCl, 10% Glycerol) and fractions were separated by SDS/PAGE and Coomassie stained to verify purified His_6_-SUMO-FliW^WT/N108Y/K123E/Q124R^ protein. Purified His_6_-SUMO-FliW^WT/N108Y/K123E/Q124R^ protein was combined with ubiquitin ligase (protease) and cleavage buffer and incubated overnight at 4 °C to cleave the SUMO tag (REF). The cleavage reaction was combined with Ni-NTA beads, incubated for 2 h at 4 °C and centrifuged to pellet the resin. Removal of the SUMO tag was verified by SDS/PAGE and Coomassie staining. Supernatant for purified FliW^WT/N108Y/K123E/Q124R^ protein was further cleaned by size exclusion chromatography on a Superdex 75 16/60 (GE Healthcare) column using CsrA gel filtration buffer (20 mM Tris-HCl pH 8.0, 200 mM NaCl, 10% Glycerol) and fractions were separated by SDS/PAGE and Coomassie stained to verify purified FliW^WT/N108Y/K123E/Q124R^ protein. FliW^WT/N108Y/K123E/Q124R^ proteins were stored at −80 °C. Concentration of FliW^WT/N108Y/K123E/Q124R^ was determined by Bradford assay (Biorad).

### Western blotting

*B. subtilis* strains were grown in LB to OD_600_ ~1.0, 1 ml sample was harvested by centrifugation, and resuspended to 10 OD_600_ in Lysis buffer (20 mM Tris pH 7.0, 10 mM EDTA, 1 mg/ml lysozyme, 10 μg/ml DNAse I, 100 μg/ml RNAse I, 1 mM PMSF) and incubated 60 minutes at 37°C. 10 μl of lysate was mixed with 2 μl 6x SDS loading dye. Samples were separated by 15% Sodium dodecyl sulfate-polyacrylamide gel electrophoresis (SDS-PAGE). The proteins were electroblotted onto nitrocellulose and developed with either anti-FliW (1:20,000) (Mukherjee et al, 2016) or anti-SigA (1:80,000; generous gift of Masaya Fujita, University of Houston) and a 1:10,000 dilution secondary antibody (horseradish peroxidase-conjugated goat anti-rabbit immunoglobulin G). Immunoblot was developed using the Immun-Star HRP developer kit (Bio-Rad).

### *In vivo* cross-linking

*B. subtilis* was grown to mid-log phase at 37°C in LB broth supplemented with 0.1mM IPTG. 10 mL culture was harvested by centrifugation, and resuspended in 10 mL of pH 7.4 PBS (137 mM NaCl, 2.7 mM KCl, 10 mM Na_2_HPO_4_, and 2 mM KH_2_PO_4_). Formaldehyde (0.3% final concentration) was added to samples and rocked at room temperature for 1 h. Next, 0.75 ml 2M glycine was added to samples and rocked at room temperature for 10 min to quench the crosslinker. Samples were centrifuged and resuspended to 50 OD_600_ in lysis buffer [20 mM Tris-HCl (pH 7.0), 10 mM EDTA, 1 mg ml^−1^ lysozyme, 10 μg ml^−1^ DNase I, and 1 mM PMSF] and incubated at 37°C for 1 h. Lysed samples were diluted 1:10 for samples probed with α–FliW antibody. 6X SDS loading dye was added and incubated for 15 min at room temperature. Samples were assayed following the Western blot protocol above. Blots were probed with either α–CsrA (1:10,000; Mukherkee et al., 2016) or α–FliW (1:20,000; Mukherjee et al., 2016) antibodies.

### GST-CsrA^WT/D41K^ and FliW^WT/mutant^ interaction pull-down assay

Glutathione-Sepharose beads were washed with T(0.1) buffer [25 mM Tris-HCl (pH 8.0), 20% glycerol, 100 mM NaCl, 1 mM DTT, 1X Protease inhibitor cocktail (from Roche), 1 mg/mL Bovine Serum Albumin (from Sigma)]. Sixty μl of washed beads was mixed with 60 μl of 0.25 μM GST-CsrA^WT/D41K^ protein and rotated on labquake at 4°C for 2 h. Next the beads bound to GST-CsrA^WT/D41K^ protein were centrifuged at 1000 rpm for 2 min and the pellet was washed twice with T(1.0) [25 mM Tris-HCl (pH 8.0), 20% glycerol, 1M NaCl, 1 mM DTT, 1X Protease inhibitor cocktail (Roche), 1 mg/mL Bovine Serum Albumin (from Sigma)] and again twice with T(0.1). Then 60 μl of increasing concentrations of FliW^WT/N108Y/K123E/Q124R^ (0.125, 0.167, 0.25, 0.333, and 0.5 μM) were added to the washed beads bound to GST-CsrA^WT/D41K^ and rotated on labquake at 4°C for 2 hours. The samples were centrifuged at 1000 rpm for 2 min and 40 μl of the supernatant was saved. The pellet was washed 4 times with T(0.1). The supernatant and pellet fractions were subjected to SDS-PAGE analysis and Western blot analysis following the protocol above. Blots were probed with either α–CsrA (1:40,000; Mukherjee et al., 2016) or α–FliW (1:60,000; Mukherjee et al., 2016) antibodies.

### Structural analysis

The FliW-CsrA 3-dimensional structure (PDB 5DMB; Altegoer et al., 2016) was analyzed using Chimera version 1.13.1.

## Acknowledgements

We thank Ayushi Mishra for strain construction. We thank Caroline Dunn and Stephen Olney for assistance with screening. We thank Masaya Fujita for the anti-SigA antibody. The work was funded by the National Institutes of Health R35 grant GM131783 to DBK.

---

**Figure S1.**
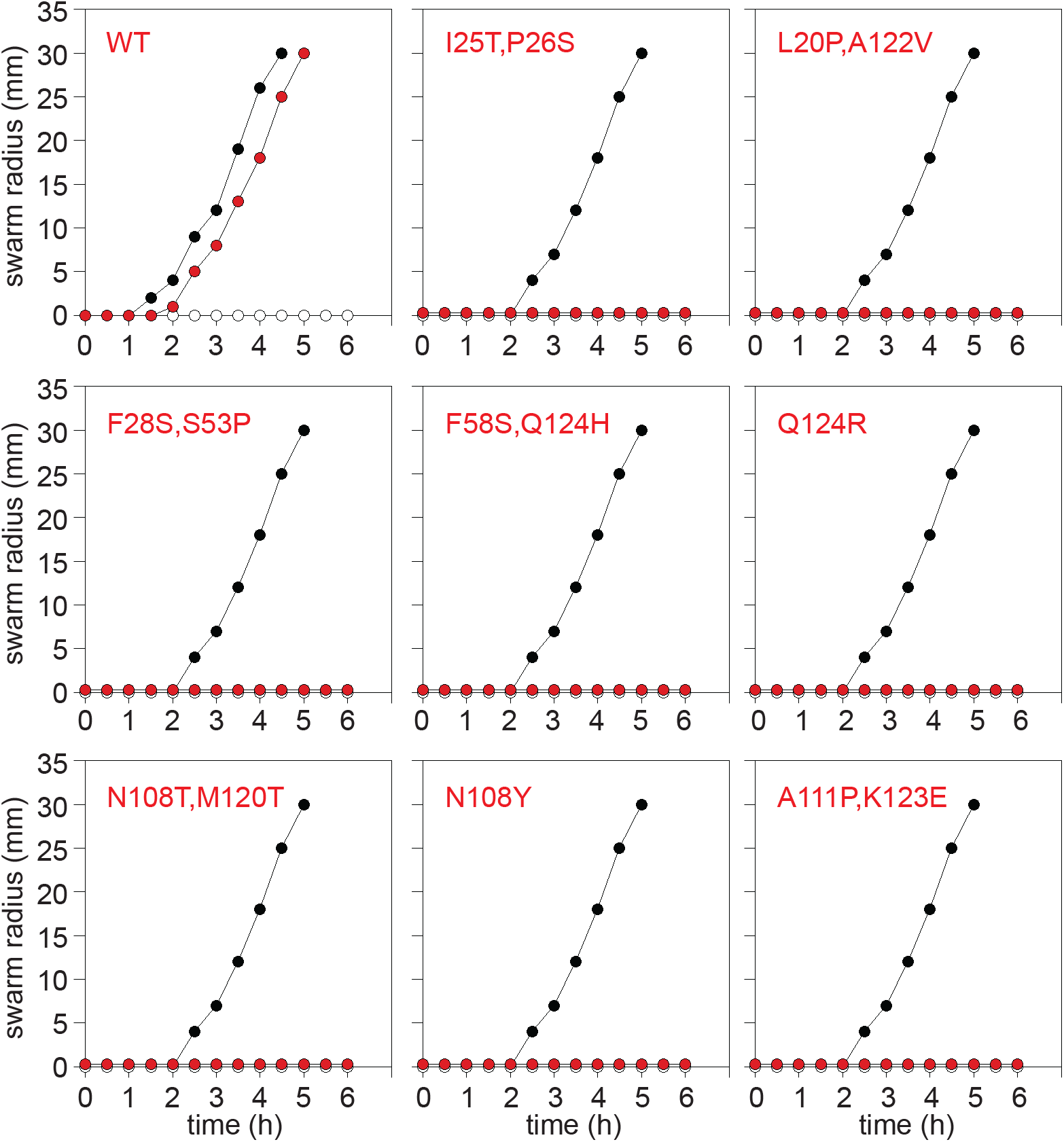
Initial loss-of-function mutations from *fliW* screen abolished motility. Quantitative swarm expansion assays of artificially induced *fliW* alleles. Each point is the average of three replicates. FliW expression was induced throughout growth and swarming by the addition of 1mM IPTG. Black circles represent WT (DK1042) and white circles represent *fliW* (DK3237) in all graphs. Red circles represent the corresponding *fliW* allele indicated in the top left corner and the following strains: *fliW amyE::P*_*IPTG*_-*fliW* (DK2371), *fliW amyE::P*_*IPTG*_-*fliW*^*I25T,P26S*^ (DK7742), *fliW amyE::P*_*IPTG*_-*fliW*^*L20P,A122V*^ (DK7387), *fliW amyE::P*_*IPTG*_-*fliW*^*F28S,S53P*^ (DK6426), *fliW amyE::P*_*IPTG*_-*fliW*^*F58S,Q124H*^ (DK7386), *fliW amyE::P*_*IPTG*_-*fliW*^*Q124R*^ (DK7389), *fliW amyE::P*_*IPTG*_-*fliW*^*N108T,M120T*^ (DK6428), *fliW amyE::P*_*IPTG*_-*fliW*^*N108Y*^ (DK6425), and *fliW amyE::P*_*IPTG*_-*fliW*^*A111P,K123E*^ (DK7741).

**Figure S2.**
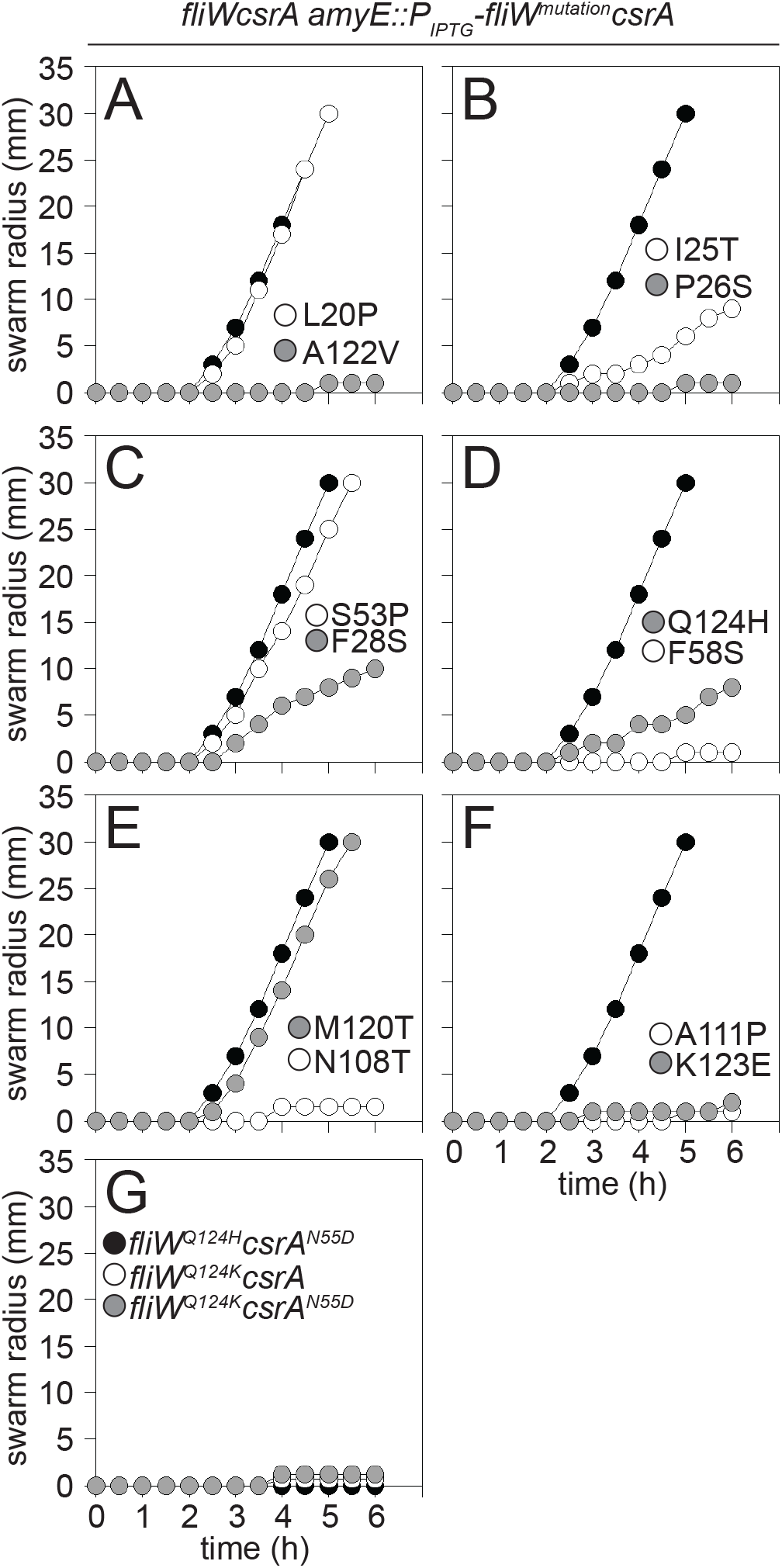
Single *fliW* mutations have differing effects on motility. A-G) Quantitative swarm expansion assays of artificially induced *fliW* alleles. Each point is the average of three replicates. FliW expression was induced throughout growth and swarming by the addition of 1mM IPTG. A-F) Black circles represent *fliWcsrA* (DK2665) in all graphs. The following strains were used to generate the panels: A) *fliWcsrA amyE::P*_*IPTG*_-*fliW*^*L20P*^*csrA* (DK8236, white circles) and *fliWcsrA amyE::P*_*IPTG*_-*fliW*^*A122V*^*csrA* (DK8241, grey circles). B) *fliWcsrA amyE::P*_*IPTG*_-*fliW*^*I25T*^*csrA* (DK8239, white circles) and *fliWcsrA amyE::P*_*IPTG*_-*fliW*^*P26S*^*csrA* (DK8240, grey circles). C) *fliWcsrA amyE::P*_*IPTG*_-*fliW*^*F28S*^*csrA* (DK8244, grey circles) and *fliWcsrA amyE::P*_*IPTG*_-*fliW*^*S53P*^*csrA* (DK8243, white circles). D) *fliWcsrA amyE::P*_*IPTG*_-*fliW*^*Q124H*^*csrA* (DK8238, grey circles) and *fliWcsrA amyE::P*_*IPTG*_-*fliW*^*F58S*^*csrA* (DK8242, white circles). E) *fliWcsrA amyE::P*_*IPTG*_-*fliW*^*M120T*^*csrA* (DK8246, grey circles) and *fliWcsrA amyE::P*_*IPTG*_-*fliW*^*N108T*^*csrA* (DK8245, white circles). F) *fliWcsrA amyE::P*_*IPTG*_-*fliW*^*A111P*^*csrA* (DK8237, white circles) and *fliWcsrA amyE::P*_*IPTG*_-*fliW*^*K123E*^*csrA* (DK7762, grey circles). G) *fliWcsrA amyE::P*_*IPTG*_-*fliW*^*Q124H*^*csrA*^*N55D*^ (DK8380, black circles), *fliWcsrA amyE::P*_*IPTG*_-*fliW*^*Q124K*^*csrA* (DK8346, white circles), and *fliWcsrA amyE::P*_*IPTG*_-*fliW*^*Q124K*^*csrA*^*N55D*^ (DK8347, grey circles).

**Figure S3.**
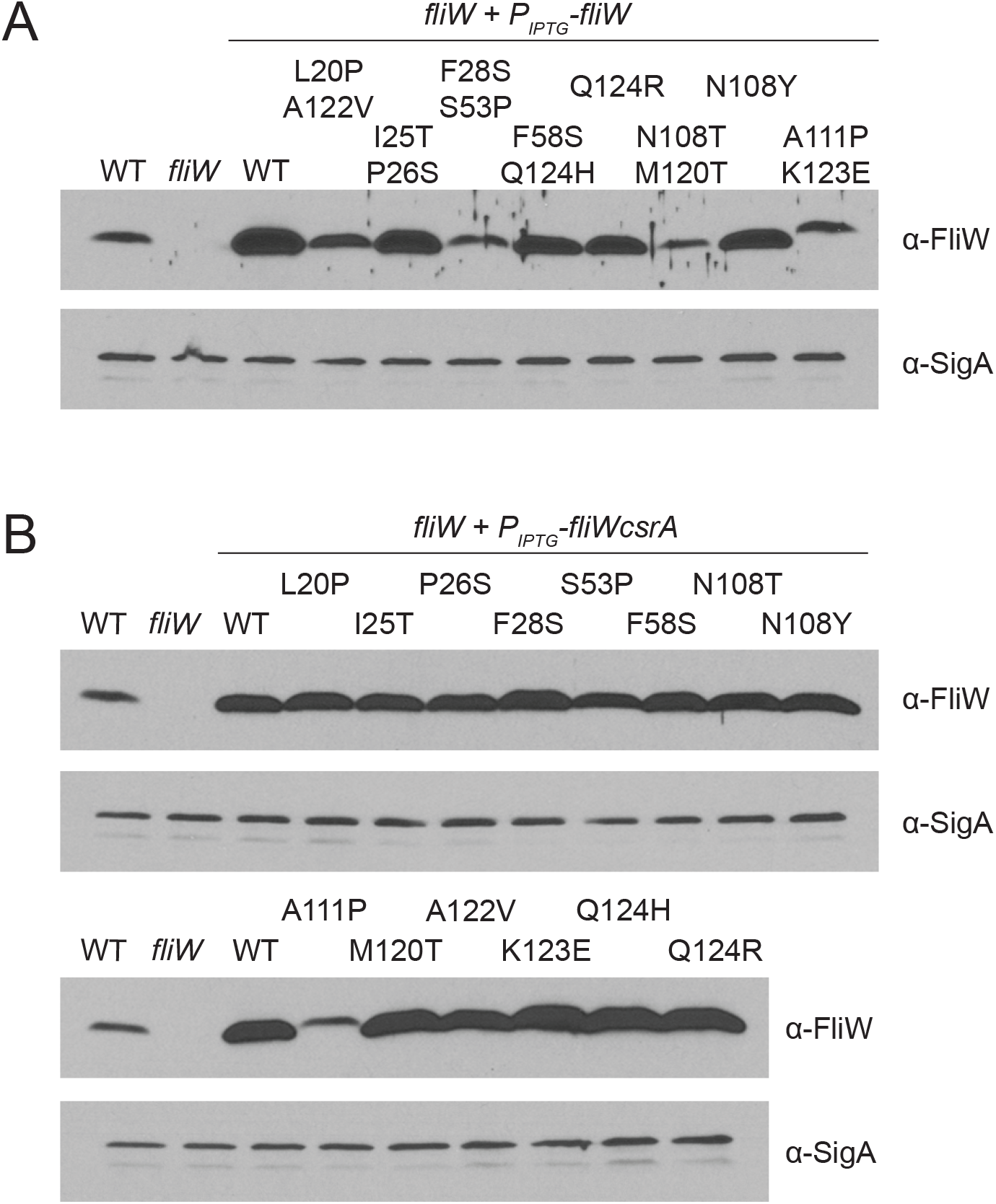
Protein stability of initial loss-of-function FliW alleles and separated single FliW mutations identified in screen. A-C) Western blot analysis of *B. subtilis* cells grown to mid-log in the presence of 1 mM IPTG to induce expression of *fliW* alleles. Whole-cell lysates were probed separately with primary antibodies raised against FliW and SigA. SigA is used as a loading control. The following strains were used to generate the panels: A) Wild type (DK1042), *fliW* (DK3237), *fliW amyE::P*_*IPTG*_-*fliW* (DK2371), *fliW amyE::P*_*IPTG*_-*fliW*^*L20P,A122V*^ (DK7387), *fliW amyE::P*_*IPTG*_-*fliW*^*I25T,P26S*^ (DK7742), *fliW amyE::P*_*IPTG*_-*fliW*^*F28S,S53P*^ (DK6426), *fliW amyE::P*_*IPTG*_-*fliW*^*F58S,Q124H*^ (DK7386), *fliW amyE::P*_*IPTG*_-*fliW*^*Q124R*^ (DK7389), *fliW amyE::P*_*IPTG*_-*fliW*^*N108T,M120T*^ (DK6428), *fliW amyE::P*_*IPTG*_-*fliW*^*N108Y*^ (DK6425), and *fliW amyE::P*_*IPTG*_-*fliW*^*A111P,K123E*^ (DK7741). B) Top. WT (DK1042), *fliWcsrA* (DK2665), *fliWcsrA amyE::P*_*IPTG*_-*fliWcsrA* (DK7567), *fliWcsrA amyE::P*_*IPTG*_-*fliW*^*L20P*^*csrA* (DK8236), *fliWcsrA amyE::P*_*IPTG*_-*fliW*^*I25T*^*csrA* (DK8239), *fliWcsrA amyE::P*_*IPTG*_-*fliW*^*P26S*^*csrA* (DK8240), *fliWcsrA amyE::P*_*IPTG*_-*fliW*^*F28S*^*csrA* (DK8244), *fliWcsrA amyE::P*_*IPTG*_-*fliW*^*S53P*^*csrA* (DK8243), *fliWcsrA amyE::P*_*IPTG*_-*fliW*^*F58S*^*csrA* (DK8242), *fliWcsrA amyE::P*_*IPTG*_-*fliW*^*N108T*^*csrA* (DK8245), and *fliWcsrA amyE::P*_*IPTG*_-*fliW*^*N108Y*^*csrA* (DK8326). Bottom. Wild type (DK1042), *fliWcsrA* (DK2665), *fliWcsrA amyE::P*_*IPTG*_-*fliWcsrA* (DK7567), *fliWcsrA amyE::P*_*IPTG*_-*fliW*^*A111P*^*csrA* (DK8237), *fliWcsrA amyE::P*_*IPTG*_-*fliW*^*M120T*^*csrA* (DK8246), *fliWcsrA amyE::P*_*IPTG*_-*fliW*^*A122V*^*csrA* (DK8241), *fliWcsrA amyE::P*_*IPTG*_-*fliW*^*K123E*^*csrA* (DK7762), *fliWcsrA amyE::P*_*IPTG*_-*fliW*^*Q124H*^*csrA* (DK8238), and *fliWcsrA amyE::P*_*IPTG*_-*fliW*^*Q124R*^*csrA* (DK8325).

**Figure S4.**
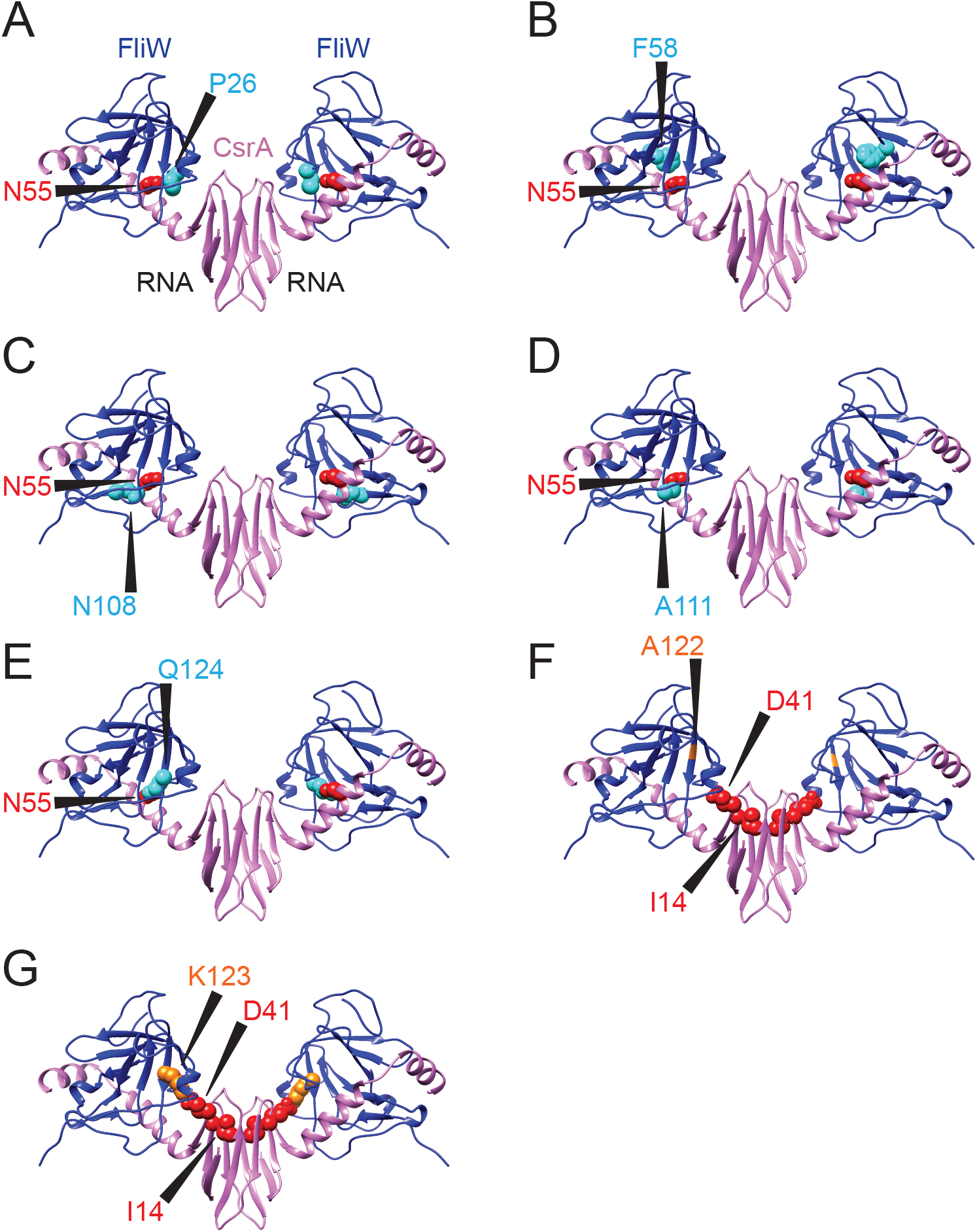
Individual residues altered by FliW loss-of-function mutations depicted on the FliW-CsrA complex. A-G) Three dimensional structures of the FliW-CsrA complex from *Geobacillus thermodenitrificans* (PDB 5DMB, REF) projected from the side. The FliW proteins are colored blue and the CsrA dimer is colored lavender. “RNA” indicates the region in which RNA would otherwise be bound. A-E) Location of each individual residue constituting the “CsrA^N55^ proximal cluster”. FliW residues within the cluster that were mutated, A) P26, B) F58, C) N108, D) A111, and E) Q124, are shown in space fill, colored cyan, and indicated by a caret. CsrA residue N55 is shown in space fill and colored red. G-H) Location of each individual residue constituting the “CsrA core proximal cluster”. FliW residues within the cluster that were mutated, G) A122 and H) K123 are shown in space fill, colored orange, and indicated by a caret. Note that FliW residue 122 of the *G. thermodenitrificans* is a glycine and not an alanine. CsrA residue D41 and I14 are shown in space fill and colored red.

**Figure S5.**
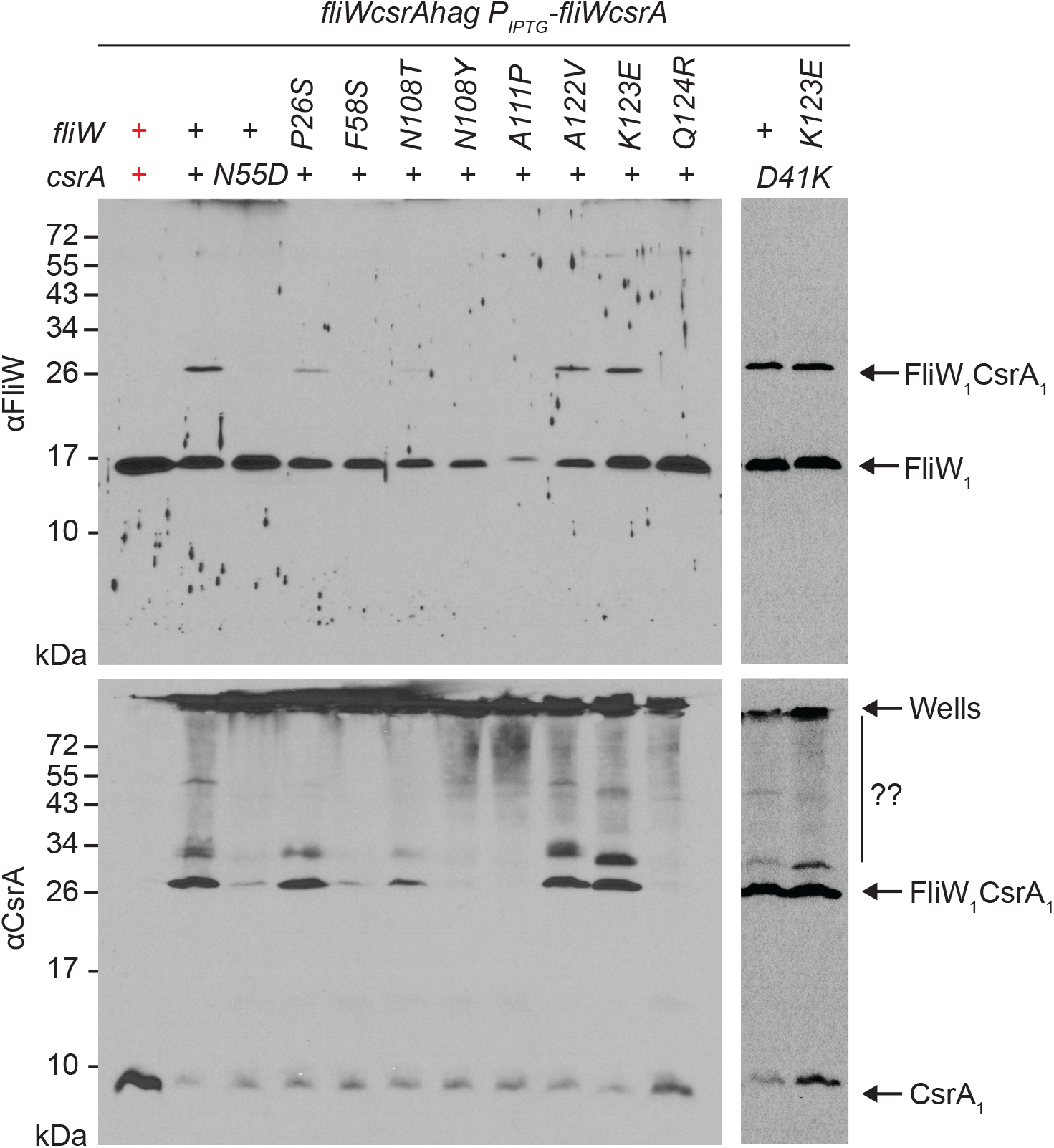
Mutation of FliW “CsrA^N55^ proximal” residues abolish CsrA interaction in vivo but mutation of FliW “CsrA core proximal” residues do not. Whole-cell lysates were grown in the presence of 0.1 mM IPTG, crosslinked with (black text) or without (red text) 0.3% formaldehyde, and subjected to Western blot analysis separately using primary antibodies against FliW (Top) and CsrA (Bottom). Samples were prepared from strains containing a triple deletion of *fliW*, *csrA*, and *hag* at their respective native sites (*fliWcsrAhag*). The introduction of the *hag* deletion was to abolish FliW-Hag interaction and reduce the complexity of the interaction species. Translational coupling of *fliW* and *csrA* was maintained by co-expressing both proteins from the same IPTG-inducible promoter integrated at an ectopic site (*amyE::P*_*IPTG*_-*fliWcsrA*). The *csrA* and *fliW* allele is expressed at the top of the panel using either “+” for wild type or the substitution as indicated. The location of CsrA monomer (CsrA_1_), FliW monomer (FliW_1_) and FliW-CsrA monomer complex (FliW_1_CsrA_1_) are indicated by carets. CsrA dimer (CsrA_2_) is not observed, likely because the interaction is too tight to permit formaldehyde crosslinking in this analysis. Additional species (??) are observed in cells expressing FliW alleles that maintain some level of binding to CsrA but the molecular nature of this species is unknown. “Wells” indicates the location of high signal that was not resolved in the gel. The following strains were used to generate lysates for both panels: Top and bottom left) *fliWcsrAhag amyE::P*_*IPTG*_-*fliWcsrA* (DK7974), *fliWcsrAhag amyE::P*_*IPTG*_-*fliWcsrA*^*N55D*^ (DK8303), *fliWcsrAhag amyE::P*_*IPTG*_-*fliW*^*P26S*^*csrA* (DK8282), *fliWcsrAhag amyE::P*_*IPTG*_-*fliW*^*F58S*^*csrA* (DK8284), *fliWcsrAhag amyE::P*_*IPTG*_-*fliW*^*N108T*^*csrA* (DK8304), *fliWcsrAhag amyE::P*_*IPTG*_-*fliW*^*N108Y*^*csrA* (DK8113), *fliWcsrAhag amyE::P*_*IPTG*_-*fliW*^*A111P*^*csrA* (DK8330), *fliWcsrAhag amyE::P*_*IPTG*_-*fliW*^*A122V*^*csrA* (DK8241), *fliWcsrAhag amyE::P*_*IPTG*_-*fliW*^*K123E*^*csrA* (DK8286), and *fliWcsrAhag amyE::P*_*IPTG*_-*fliW*^*Q124R*^*csrA* (DK8108). Top and bottom right) *fliWcsrAhag amyE::P*_*IPTG*_-*fliWcsrA*^*D41K*^ (DK8379) and *fliWcsrAhag amyE::P*_*IPTG*_-*fliW*^*K123E*^*csrA*^*D41K*^ (DK8378).

**Figure S6.**
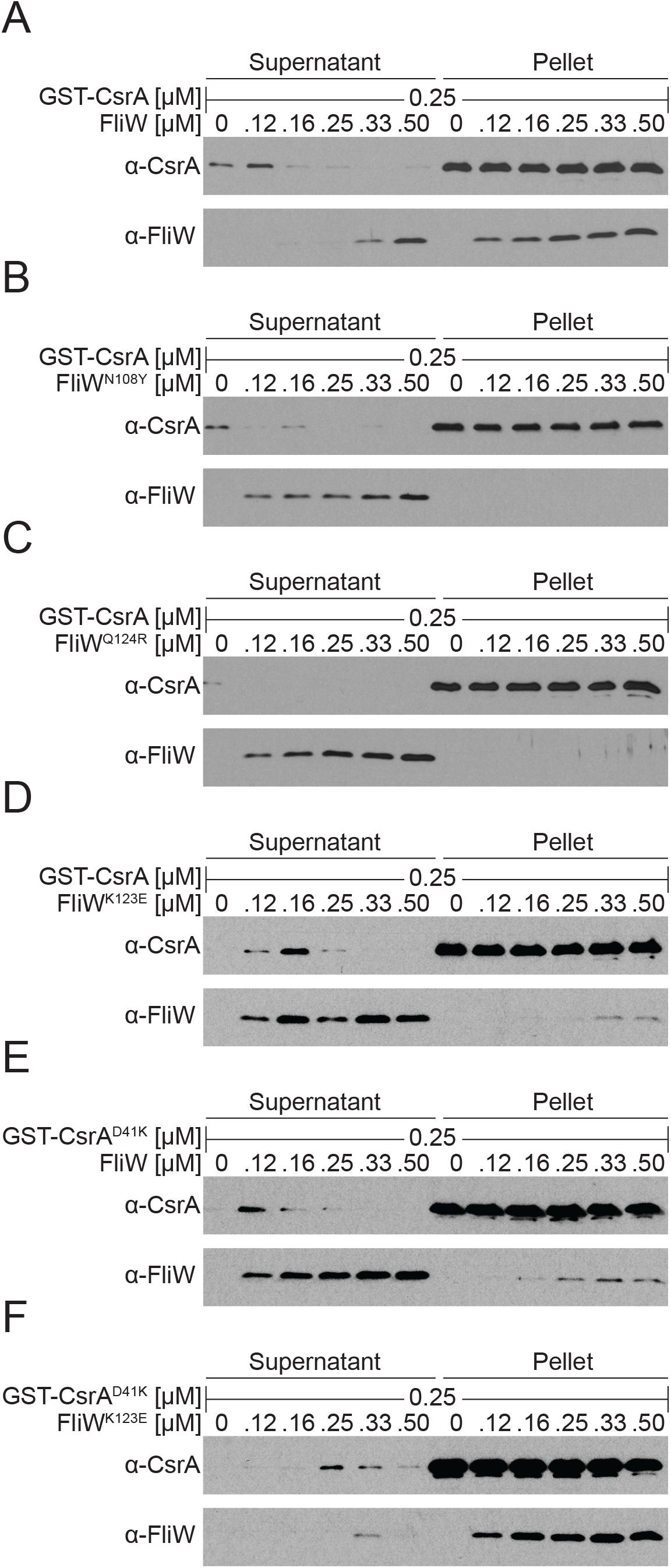
Differential binding of *lof* FliW alleles to CsrA^WT^ and CsrA^D41K^*in vitro*. A-F) A protein pull-down assay using purified GST-CsrA^WT^ (A-D) or GST-CsrA^D41K^ (E-F) proteins at the indicated amount was loaded onto a glutathione-sepharose column and incubated with the indicated amounts of purified FliW^WT^ (A and E), FliW^N108Y^ (B), FliW^Q124R^ (C), or FliW^K123E^ (D and F) protein in the presence of BSA. Samples were subjected to Western blot analysis and probed with primary antibodies to FliW and CsrA. “Supernatant” indicates the proteins that failed to bind to the beads, and “pellet” indicates the proteins that remained bound to the beads following a series of washes. Note, that in some instances CsrA can be seen in the supernatant lanes and this is due to resin being accidentally collected while taking the supernatant sample.

**Table S1.**
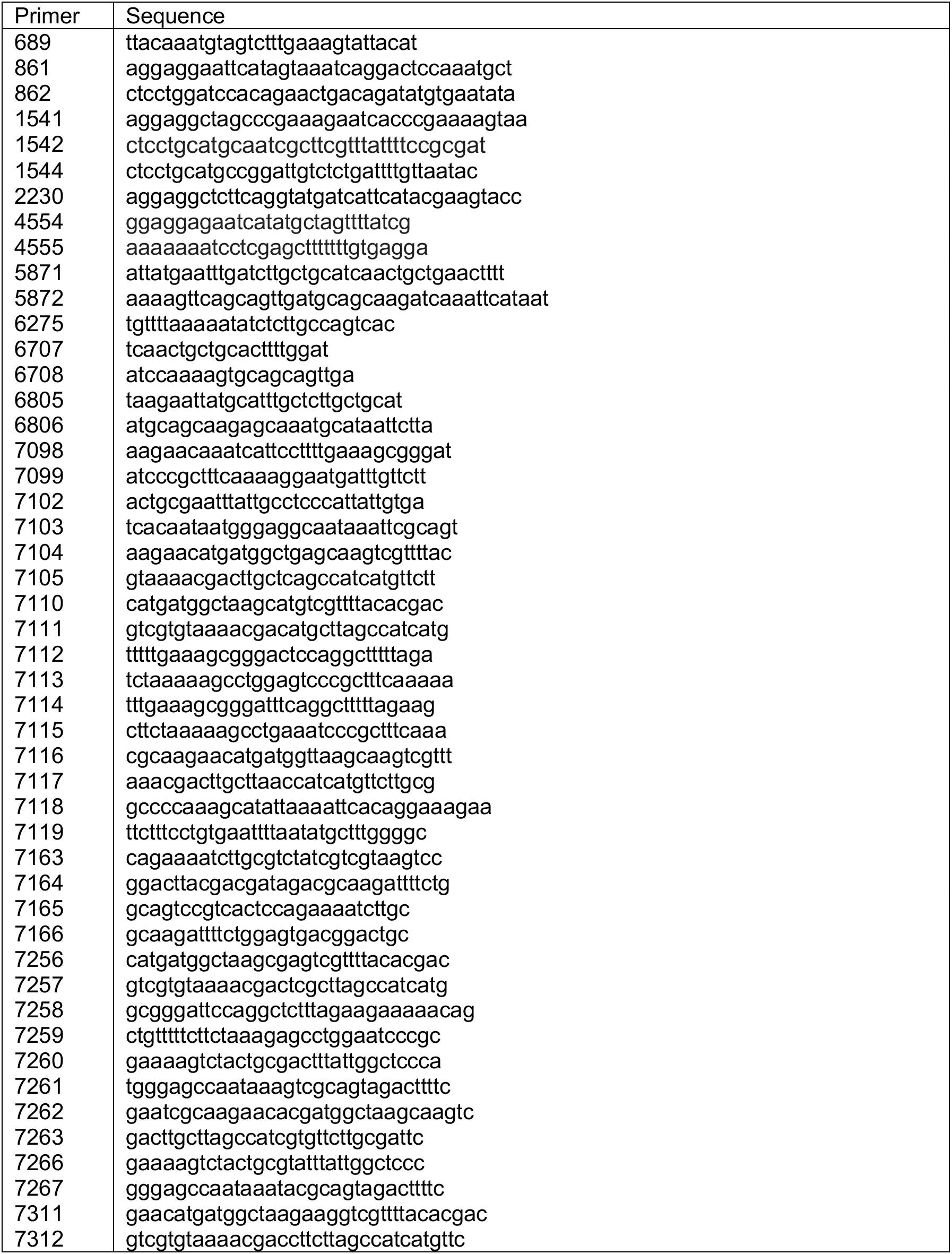
Primers

**Table S2.**
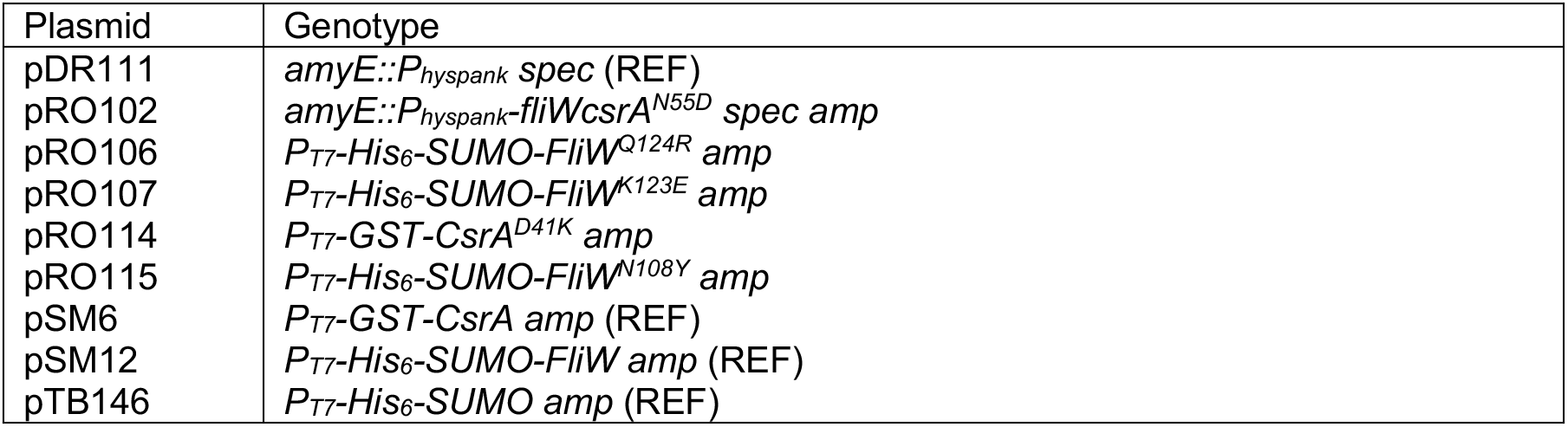
Plasmids

